# Laws of diversity and variation in microbial communities

**DOI:** 10.1101/680454

**Authors:** Jacopo Grilli

**Affiliations:** Santa Fe Institute, 1399 Hyde Park Road, Santa Fe, NM 87501, USA; The Abdus Salam International Centre for Theoretical Physics (ICTP), Strada Costiera 11, 34014 Trieste, Italy

## Abstract

How coexistence of many species is maintained is a fundamental and unanswered question in ecology. Coexistence is a puzzle because we lack a quantitative understanding of the variation in species presence and abundance. Whether variation in ecological communities is driven by deterministic or random processes is one of the most controversial issues in ecology. Here, we study the variation of species presence and abundance in microbial communities from a macroecological standpoint. We identify three novel, fundamental, and universal macroecological laws that characterize the fluctuation of species abundance across communities and over time. These three laws — in addition to predicting the presence and absence of species, diversity and other commonly studied macroecological patterns — allow to test mechanistic models and general theories aiming at describing the fundamental processes shaping microbial community composition and dynamics. We show that a mathematical model based on environmental stochasticity quantitatively predicts the three macroecological laws, as well as non-stationary properties of community dynamics.

---

No two ecological communities are alike, as species composition and abundance vary widely. Understanding the main forces shaping the observed variation is a fundamental goal of ecology, as it connects to multiple issues, from the origin of species coexistence to control and conservation. Thanks to the fast and recent growth of data availability for microbial communities, we have a detailed understanding of which environmental factors affect community variability [1–4] and, sometimes, the genetic drivers determining the response to different environmental conditions [5, 6]. This qualitative understanding of the correlates, and potential causes, of the observed variation does not parallel with a quantitative understanding of its fundamental and general properties [7–9]. We do not, in fact, quantitatively understand how species presence and abundance fluctuate across microbial communities, and which ecological forces are responsible for these fluctuations. Macroecology, the study of ecological communities through patterns of abundance, diversity, and distribution [10], is a promising approach to study quantitatively variation in microbial communities [11–13]. Here we show that three novel macroecological laws describe the fluctuations of abundance and diversity. These three ecological laws are universal, since they hold across biomes and for both cross-sectional and longitudinal data, and are fundamental, as they suffice to predict, fitting no parameters, the scaling of diversity and other commonly studied macroecological patterns, such as the Species Abundance Distribution. Macroecological patterns are the bridges from uncharacterized variation to ecological processes and mechanisms. We show that a model based on environmental stochasticity can reproduce the three macroecological laws, as well as dynamic patterns in longitudinal data. Both data and model show that, at the taxonomic resolution commonly used, competitive exclusion is rare and variation of species presence and abundance is mostly due to environmental fluctuations. Our results are a solid basis for inferring and modeling biotic and abiotic interactions in a rigorous data-driven quantitative way.

The most studied pattern in ecology is the Species Abundance Distribution (SAD) [14, 15], which is defined as the distribution of abundances across species in a community. Multiple functional forms, and consequently multiple models, have been proposed to describe the empirical SAD in microbial communities [12]. While SADs are highly studied and characterized, it is often neglected that three distinct and independent sources of variation influence their shape: sampling error, fluctuation of abundances of individual species, and variability in abundance across species. We disentangle these sources of variation in three novel, more fundamental, macroecological laws.

The first pattern we consider is the Abundance Fluctuation Distribution (AFD), which is defined as the distribution of abundances of a species across communities (see Figure 1a). By properly accounting for sampling errors (see Appendix and Supplementary Section S2), we show that the Gamma distribution, with species’ dependent parameters (see Figure 1b and Supplementary Figure S1), well describes the AFD across biomes and species. Whichever ecological process is at the origin of species’ abundance variation, it manifests regularly and consistently.

**FIG. 1:**
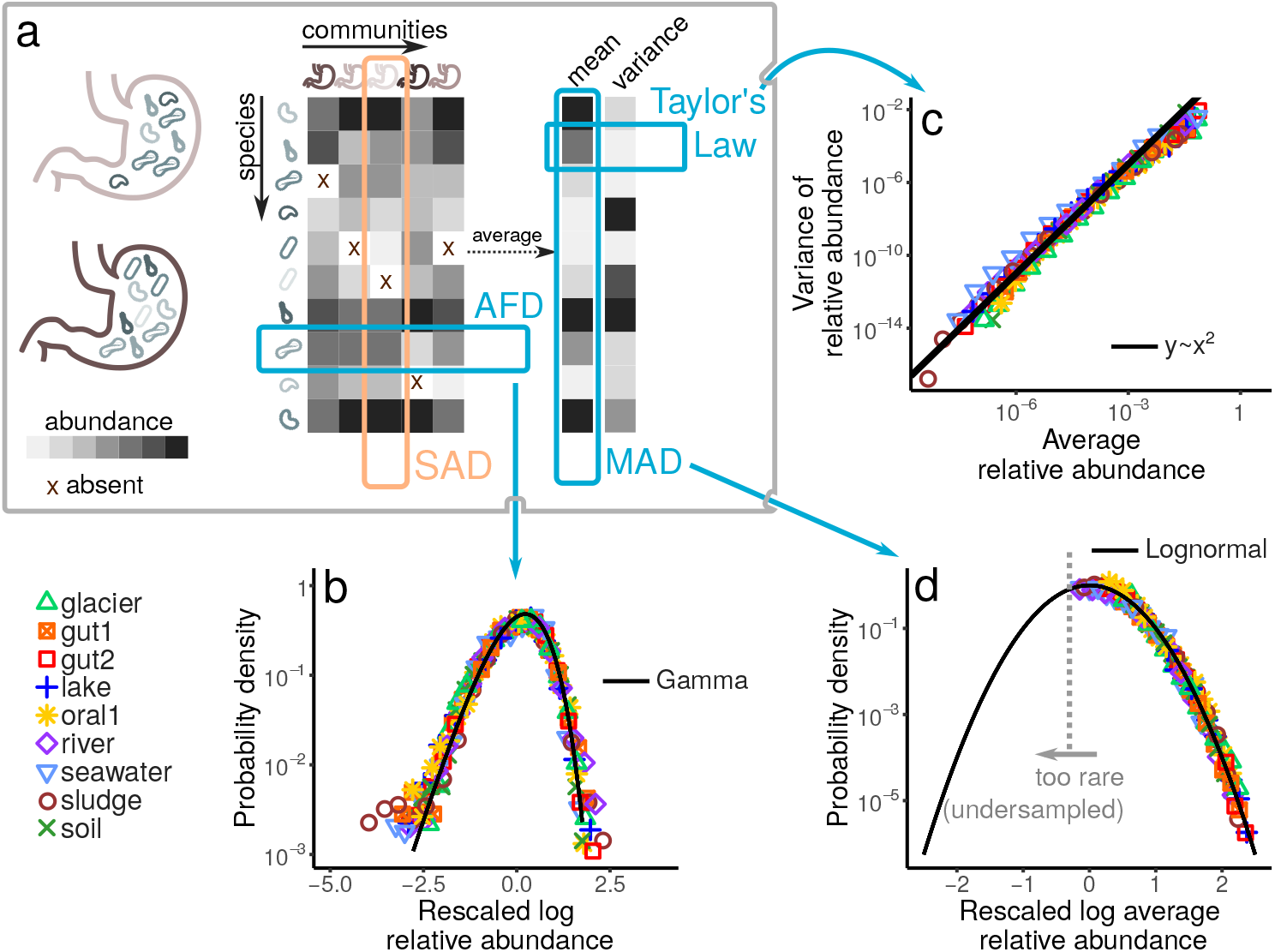
Three laws of microbial community composition. Panel a illustrates the cross-sectional data and what the three laws describe. Similarly to other component systems [31], the abundance of a given microbial species (see Appendix and Supplementary Section S1 for definitions in different datasets) in a community corresponds to the entry of a matrix where columns are communities and rows are species. One of the most commonly studied patterns in ecology is the species abundance distribution (SAD), which describes the fluctuations of abundance across species (rows) in a community (column). Instead of focusing on the SAD, we study the Abundance Fluctuation Distribution (AFD), which describes the distribution of abundances of a species across communities. We consider cross-sectional data from 7 projects and 9 biomes (colored symbols), collected and processed in different ways (see Appendix). Panel b shows that a Gamma distribution (solid black line) closely matches the AFD (see Supplementary Section S3). In real data, sampling errors strongly affect this pattern (see Supplementary Section S2). Here, we average the AFD over the species that are always present in a biome, by rescaling their log relative abundance. In Supplementary Section S2 we describe a method to disentangle the AFD from the variation introduces by sampling, showing that a Gamma distribution also describes the AFD of rarer species (see Supplementary Section S3 and Supplementary Figure S2). Since the AFD is Gamma distributed for all species, the average abundance and its variance of each species are enough to describe the fluctuations of a species across communities. Panel c shows that mean and variance are not independent across species, a relationship known as Taylor’s Law. The variance is, in fact, proportional to the square of the mean (solid line), implying that the coefficient of variation of the abundance fluctuations is constant across species. Supplementary Figure S2 describes how we removed the effect of sampling in order to obtain Taylor’s Law. Taylor’s Law (together with a Gamma AFD) implies that a single parameter per species (the average abundance) is enough to recapitulate its distribution of fluctuations. Panel d shows that the Mean Abundance Distribution (MAD), which is defined as the distribution of mean abundance (obtained by averaging over communities) across species, is Lognormally distributed (see Supplementary Section S2 B for relative abundance normalization). Colored symbols represent data (where we rescaled abundances so that the logarithm had mean zero and variance one), while the black line corresponds to a Lognormal pdf. Note that we expect sampling to strongly influence this pattern: it is less likely to observed species that are rare (left tail of the MAD, see Supplementary Section S7). By determining the parameters of the MAD using the observed data we can estimate the number of unobserved species (see Supplementary Section S7). The two parameters of the best Lognormal fit to the MAD are biome dependent (see Appendix), and, together with the total diversity and the coefficient of variation of the AFD (which is species independent), they can be used to predict other patterns of abundance and diversity.

The probability that a Gamma-distributed variable is zero vanishes. A direct consequence of the first macroecological law (a Gamma AFD) is that all instances in which a species is absent should be imputed to sampling error. We directly test this prediction in two ways. First, using Bayesian model selection, we show that a Gamma AFD is statistically superior to models including species absence (see Appendix and Supplementary Section S3). Second, if the absence is caused by sampling error, we can predict the occupancy of a species, defined as the fraction of communities where it is present, from the AFD. Figure 2 shows that we can indeed predict the occupancy from the first two moments of species abundance fluctuations. This result strongly suggests that, at the taxonomic resolution commonly used, competitive exclusion is absent or, at least, statistically irrelevant. Importantly, this result clarifies the relation between abundance and occupancy [16], which has been reported in multiple microbial systems [13, 17, 18] but has never been quantitatively characterized and explained.

**FIG. 2:**
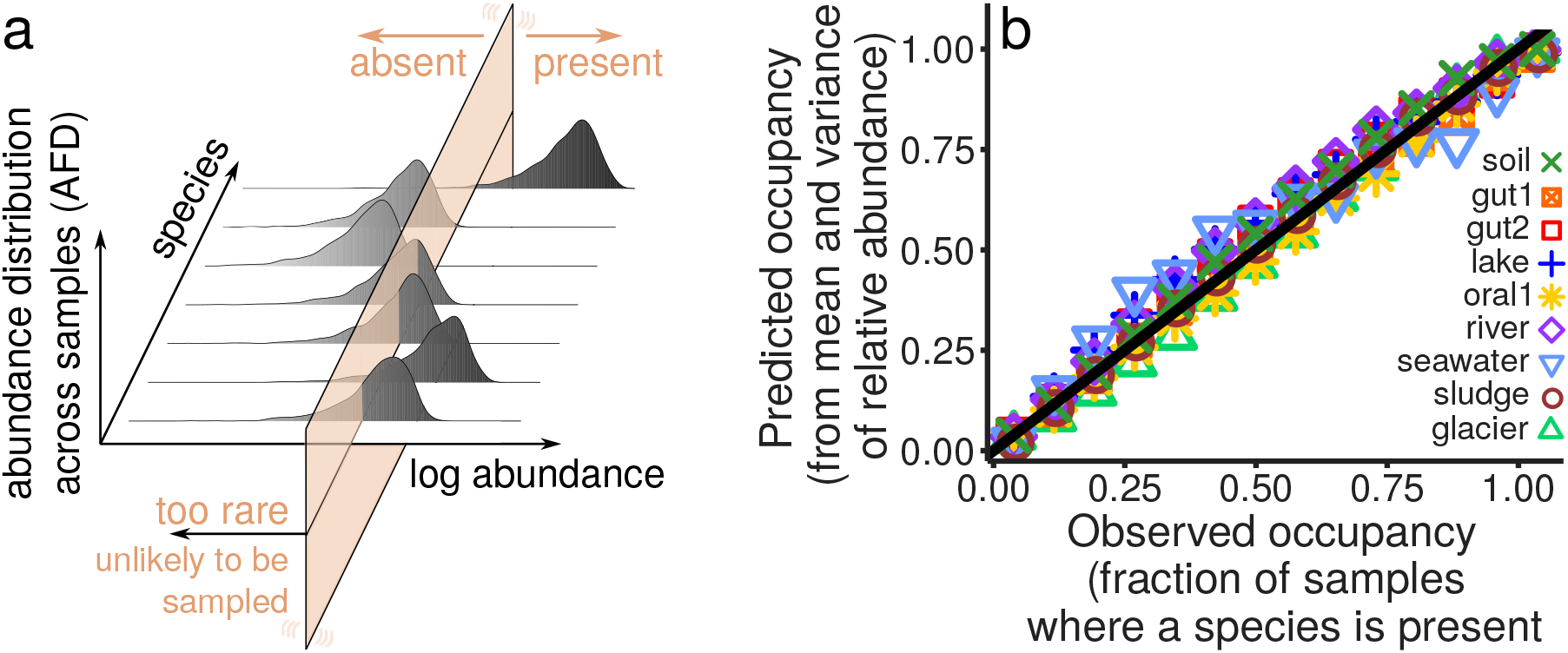
The AFD predicts presence/absence of species from fluctuations of abundance. Panel a illustrates the relationship between fluctuation in abundance and the absence of species. The fluctuations of species abundances across communities (AFD) are Gamma distributed (see Figure 1), which implies that species are absent only because of finite sampling. Panel b tests this prediction, by comparing the occupancy of species (the fraction of communities where a species is presence) in different biomes with what expected from independent sampling from Gamma distributed relative abundances (see Supplementary Section S4 and Supplementary Figure S3). By modeling explicitly sampling from Gamma AFD (which requires only the knowledge of average and variance of the abundances, see Appendix) we correctly predict species occupancy. This result implies that, at the taxonomic scale at which we are observing the community, the absent species are false negatives and therefore that there is no evidence of competitive exclusion.

The mean and variance of abundance fluctuations are therefore sufficient to characterize the full distribution of abundances of species across communities. The second macroecological law we discovered describes the relation between mean and variance of species abundance, which is often referred to as Taylor’s Law [19, 20]. Figure 1c shows that the variance scale quadratically with the mean, implying that the coefficient of variation of the abundance fluctuations is constant (with respect to mean abundance, see also Supplementary Section S5). Thanks to Taylor’s Law, we need therefore only one, instead of two, parameters per species — their average abundance — to describe species abundance fluctuations. The average species abundance is a biologically relevant quantity, as it has a reproducible dependence on the biome, and strong phylogenetic signal, with similar species having similar average abundance (see Supplementary Section S6).

Since the average abundance alone characterizes the distribution of abundance fluctuations of each species, it is natural to analyze how the average abundance is distributed across species (MAD, mean abundance distribution). Figure 1d shows that the MAD is Lognormally distributed for all the datasets considered in this work. Since in a finite number of samples rare species are likely to be never sampled, the empirical MAD displays a lower cutoff which is determined by sampling. This property allows to estimate the total diversity, under the assumption that the MAD is Lognormal also for the rarer species (see Supplementary Section S7). We find that the total diversity is typically at least twice as large as the recorded one (see Supplementary Table SS2). A Lognormal MAD also rules out neutral theory [21, 22] as an explanation of community variability. For a finite number of samples, neutral theory would, in fact, predict a Gaussian MAD (see Supplementary Section S12), which we can easily reject from the data.

The three laws presented so far — the Gamma AFD, Taylor’s Law with exponent 2 and the Lognormal MAD — can be fully parameterized for each biome knowing the first two moments of the MAD (how the mean relative abundance differs across species), the total diversity and the coefficient of variation of the AFD (what is the average variation of species’ abundance across communities, i.e., the intercept of Figure 1c). Knowing these parameters, one can generate synthetic samples for arbitrary levels of sampling and compare the statistical properties of these synthetic samples with the empirical ones. We focus on commonly studied macroecological patterns: the relation between diversity and the number of sequences sampled [23] (which is, somewhat, parallel to the Species-Area relationship [11]), the Species Abundance Distribution [14, 15], the occupancy distribution [24] and the abundance-occupancy relationship [16] (see also Supplementary Section S8 for other quantities). It is important to note that these patterns are all affected by sampling, species abundance fluctuations, and species differences. Knowing the three macroecological laws allows us to analytically calculate a prediction for these quantities (see Appendix and Supplementary Section S8). Figure 3 shows that the predictions of these macroecological patterns match the data accurately. The three laws are therefore not only universal (i.e., valid across biomes), but also fundamental: we can use them to predict other macroecological quantities.

**FIG. 3:**
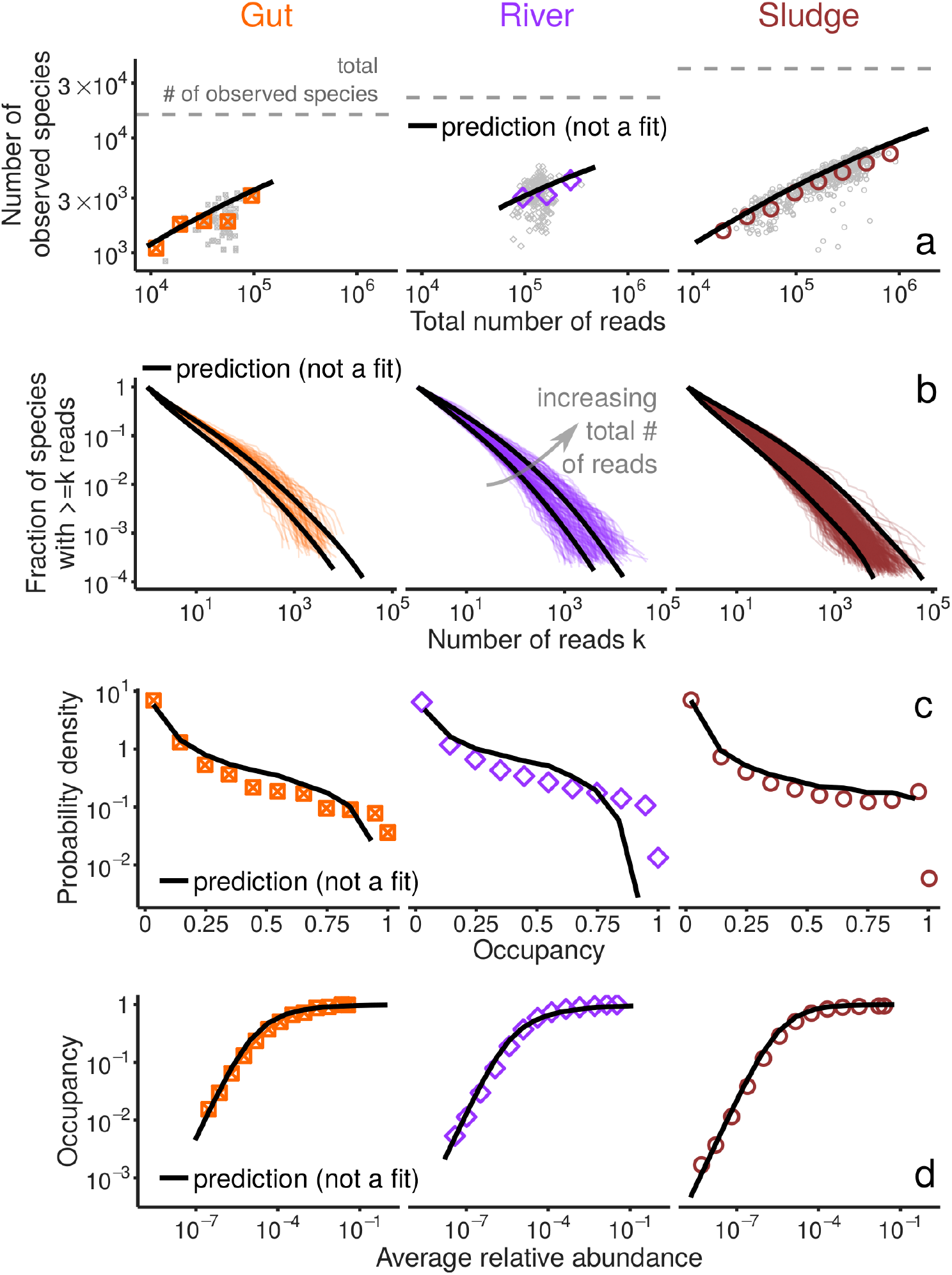
The AFD, Taylor’s Law and MAD predict quantitatively macroecological patterns. Panel a shows the scaling diversity (measured as the number of species) with the total number of sampled sequences (reads). Each gray point represents a community (colored points are averages). The black solid lines is the prediction from the three fundamental patterns and sampling (see Appendix and Supplementary Figure S9), which correctly reproduces the empirical trend. The gray dashed line corresponds to the total number of different species observed in at least one community. Panel b compares the cumulative SAD of different ecological communities (colored lines), with the prediction of the three patterns and sampling. The two solid black curves refer to the prediction with a total number of sequences equal to the smallest and the largest one of the empirical samples (see also Appendix and Supplementary Figure S22). Similarly, panel c compares the distribution of occupancy observed (colored symbols) and predicted by sampling from the prediction of the three laws (see Supplementary Figure S11). Panel d shows that the three macroecological laws accurately predict (black line) the abundance-occupancy relation [16] observed in the data (colored symbols, see also Supplementary FigureS21). Note that these predictions were obtained simply by measuring the parameters of the MAD, the total observed diversity (the gray dotted line in panel line) and the coefficient of variation of the AFD (the intercept of Figure 1c).

A question that naturally arises from the success of the AFD, together with the other two macroecological laws, in predicting the scaling of abundance and diversity translates into an ecological prediction on the nature of stochasticity. Which ecological process is responsible for the fluctuations of species abundance across communities? The ability of a Gamma AFD in predicting occupancy from its first two central moments, as illustrated in Figure 2, rules out mechanisms that explain variation as a consequence of alternative stable states driven by biotic or abiotic interactions. These mechanisms would correspond in fact to more complicated relationships between abundance and occupancy (see Supplementary Section S12), that cannot be described by a Gamma AFD. An alternative is that the variation in abundances is the effect of a mechanism with some intrinsic variability. This variability could be due to heterogeneity (e.g., two communities are different because the environmental conditions were, are and will be different) or stochasticity (e.g., two communities are different because the environmental conditions are independently fluctuating over time). We tested these two scenarios using longitudinal data (see Appendix). In the former scenario, the three macroecological laws should differ between cross-sectional and longitudinal studies. While in the latter case, they should also hold when a community is followed over time. Figure 4 shows that the three macroecological laws also hold for longitudinal data, suggesting that fluctuations in abundance are mainly due to temporal stochasticity (see also Supplementary Section S9). This result does not contradict the existence of replicable differences between communities (e.g., host genetics correlates with community composition of gut microbiome [25]). We claim that most of the variation, and not all of it, is due to temporal stochasticity. We estimated, using longitudinal data, that 80% of species abundance variation is due to temporal stochasticity (see also Supplementary Section S11). The remaining 20% can be used to detect, and is likely to contain, signatures of community heterogeneity.

**FIG. 4:**
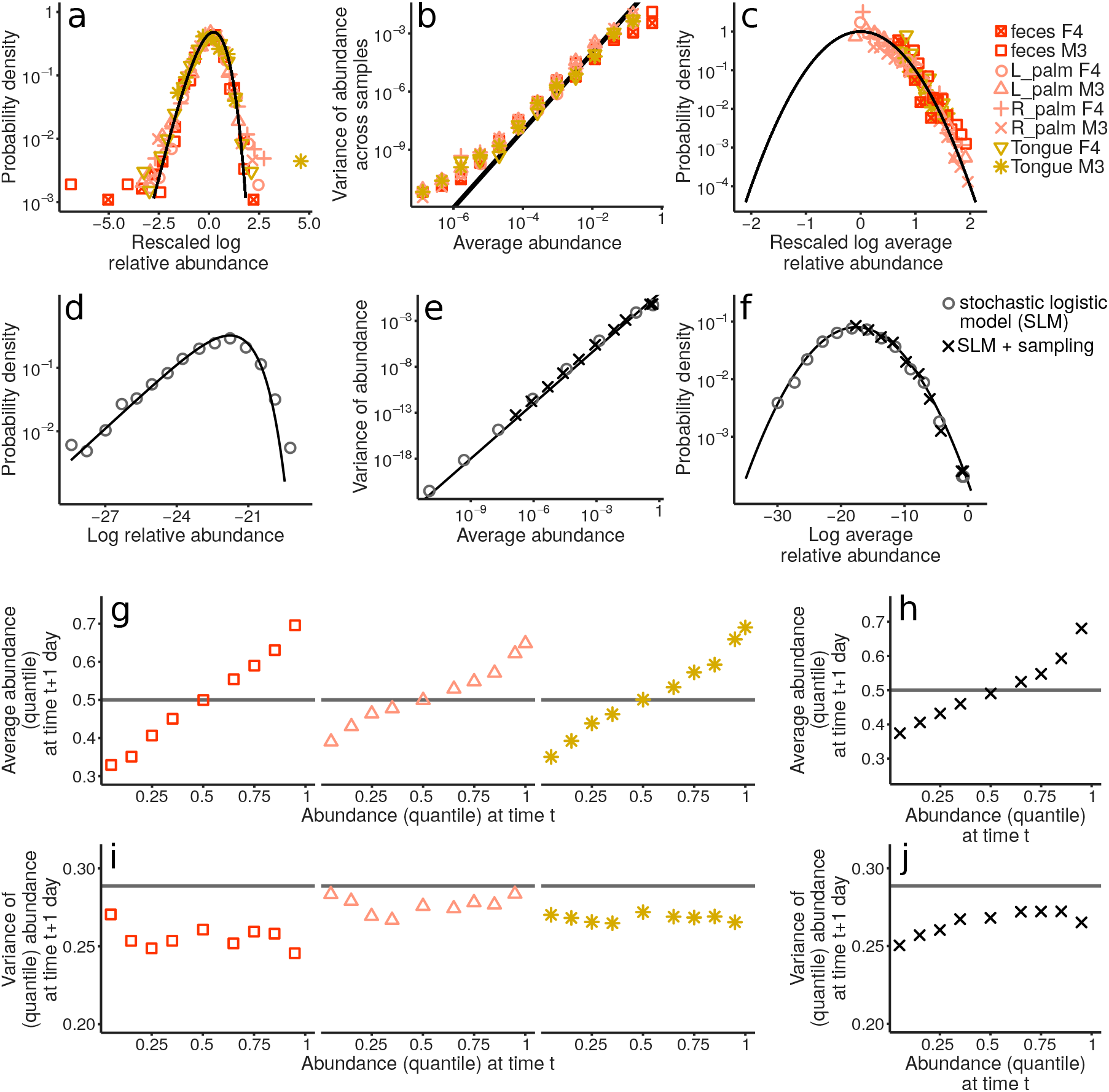
The stochastic logistic model predicts the fundamental laws in longitudinal data. Panels a,b, and c show that the AFD, the Taylor’s law, and the MAD also hold for longitudinal data (colored points, see Appendix). Panels d,e and f show that the stochastic logistic model (SLM) reproduces the empirically observed AFD, Taylor’s law and MAD, respectively. Gray circles are the results obtained with the SLM, and the black crosses the ones obtained using SLM together with sampling. Longitudinal data allow to test prediction on the dynamics. A correct model should not just be able to predict stationary properties (like the ones shown in panel a,b and c), but also non-stationary ones (i.e., transition probabilities). Panel g shows the average quantile abundance given an average quantile abundance in the previous day (averaged over species, see Appendix and Supplementary Section S9). The gray solid line shows the expected relation in the absence of time dependence. Similar to panel g, panel i shows the variance of the quantile abundance given an average quantile abundance in the previous day (averaged over species). Panels h and j show that the SLM correctly predicts the non-stationary properties shown in panels g and i (see Appendix).

The observation that variation in abundances is mostly due to stochasticity over time, together with the three macroecological laws, strongly constrains the validity of models aiming at explaining and reproducing community dynamics. We interpret stochasticity as due to environmental fluctuations (an alternative would be demographic stochasticity, see Appendix). We considered the stochastic logistic model (SLM, see Appendix) to describe species population dynamics. The SLM assumes that species populations grow logistically, with a time-dependent growth rate, which fluctuates at a faster rate than the average growth rate (i.e., the timescale associated with growth rate fluctuations is much shorter than the typical timescale of population dynamics). Figure 4 shows that the SLM can reproduce the three macroecological laws at stationarity. Note that the SLM assumes that species abundances fluctuate independently, a hypothesis that can be falsified with data (see Supplementary Section S14). Nevertheless, the SLM can be interpreted as an effective model (mean-field, in the language of statistical physics [26, 27]) capturing the statistics of individual species fluctuations.

A correct model describing population dynamics should not only reproduce the stationary distribution but also time-dependent quantities. Assuming that dynamics are Markovian, the state of the system would be fully characterized by the transition probability, which is defined as the probability of observing an abundance at time *t* + Δ*t*, conditional to the abundance at time *t*. Figure 4 shows the first two central moments of this distribution (see Appendix), for Δ*t* = 1 day. An important observation is that we can, in fact, detect a signature of dynamics: the longitudinal data, collected with a time-spacing of 1 day, display a non-trivial time correlation (with a typical relaxation time-scale equal to 19 hours, see Appendix). Figure 4 shows the SLM reproduces also the dynamics patterns, giving further validation to the hypothesis that environmental fluctuations drive the variability observed in the data.

Having a quantitative model validated with data allows in fact to extrapolate (e.g., make predictions about the future conditional to the past, or the larger scale given the smaller one), infer (i.e., measure biologically interpretable parameters) and predict (i.e., using the data one can falsify it and/or its extensions). The ability to identify a (simple) model able to quantitatively reproduce fundamental and universal, and yet non-trivial, macroecological patterns put us in the position of having a solid quantitative ground to explore the relative strength of more complicated and, perhaps, interesting ecological mechanisms. For instance, we can easily modify the SLM to include explicitly biotic or abiotic interactions. More fundamentally, while the three macroecological laws do not automatically point to any ecological mechanism (that we identified using the SLM), they clearly stem from data. Thus, they are a fact that any model aiming at quantitatively describe microbial communities cannot ignore.

## Acknowledgments

I thank S. Allesina, D. Bhat, M. Cosentino Lagomarsino, A. Kolchinsky, P. Lemos-Costa, A. Maritan, A. Mazzolini, M.A. Muñoz, M. Osella, J. Piñero, M. Sireci, R. Solé, D. Stouffer, S. Suweis, and S. Zaoli for comments and discussions at different stages of this work. Special thanks to Emilio Canzi for his inspiring ideas. J.G. was supported by an Omidyar Postdoctoral Fellowship at the Santa Fe Institute.

## Appendix

## Data

All the datasets analyzed in this work have been previously published and were obtained from EBI Metagenomics [28]. Previous publications (see Supplementary Table S1) report the original experiments and corresponding analysis. In order to test the robustness of the macroecological laws and the modeling framework presented in this work, we considered 7 datasets that differ not only for the biome considered, but also for the sequencing techniques and the pipeline used to process the data.

## Sampling and compositional data

We are interested in studying how (relative) abundance varies across communities and species. We would like to remove the effect of sampling noise, as it is not a biologically-informative source of variation. We explicitly model sampling (see Supplementary Section S2), finding that the probability of observing *n* reads of species *i* in a sample with *N* total number of reads, is given by

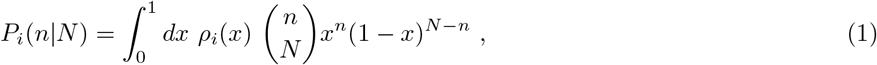

where *ρ*_*i*_(*x*) is the Abundance Fluctuation Distribution, i.e. the probability (over communities or times) that the relative abundance of *i* is equal to *x*. Note that this equation does not assume any hypothesis about independence across species or communities. It only assumes the sampling process is carried independently across communities.

Since the random variable *x*_*i*_, whose distribution is *ρ*_*i*_(*x*), is a relative abundance, we have that Σ_*i*_*x*_*i*_ = 1 (i.e., the data are compositional [29]). As discussed in Supplementary Section S2, given the range of variation of the empirical relative abundances, we can substitute eq. 1 with

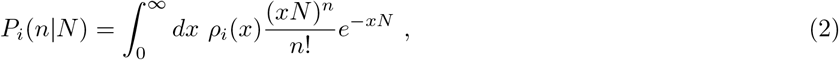

and the condition Σ_*i*_*x*_*i*_ = 1 to 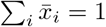, where 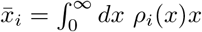 is the mean value of *x*_*i*_. Under this assumption, we can also take the limits of the integration from 0 to ∞, instead of considering them from 0 to 1.

Note that, because of sampling, the average of a function *f*(*x*) over the pdf *ρ*(*x*) differs in general from the average of *f*(*n*/*N*) over *P*(*n*|*N*)

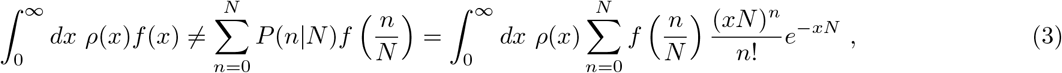

and the inequality becomes an equality only if *f*(*x*) is linear. The important difference between right- and left-hand side is often neglected in the literature. In fact, the right hand side is a good approximation of the left-hand size only in the limit *xN* ≫ 1, which is far from being realized in the data for most of the species. In Supplementary Section S2 we introduce a method to reconstruct the moments of *ρ*(*x*) from the moments of *P*(*n*|*N*). More generally, we show that it is possible to infer the moment generating function of *ρ*(*x*) from the data, which allows to reconstruct the shape of the empirical *ρ*(*x*).

## Three macroecological laws

Law #1. The Abundance Fluctuation Distribution (AFD) *ρ_i_*(·) is a Gamma distribution

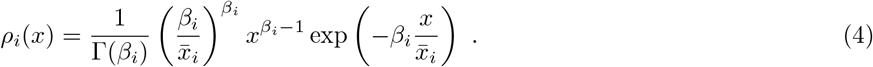

The two parameters 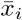 and *β*_*i*_ fully characterize the AFD of each species. The parameter *β*_*i*_ is related to the squared inverse coefficient of variation: 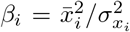, where 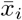 is the average abundance of species *i* and 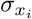 is its standard deviation. We tested this law against alternative distributions in Supplementary Section S3, obtaining a superior performance of the Gamma distribution in all the datasets considered in this study.

Law #2. The coefficient of variation of the abundance distribution is constant (does not scale with the average abundance 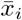). A power-law relation between mean and variance of the type 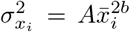 is often refereed to as Taylor’s Law [19]. In our case, it holds with *b* = 1. In particular, it implies that *β*_*i*_ = *β* for all species (see also S5).

Law #3. The average (relative) abundance 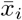 is lognormally distributed across species

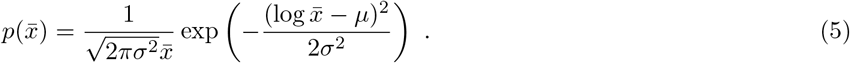

The parameter *σ* characterizes the variability in the mean abundance across species. Since we are always dealing with a finite number of (finite) samples, some species are never observed. If a species is rare enough (i.e., if 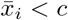, where *c* is a cutoff determined by the number of samples and the total number of reads in each sample), it becomes extremely unlikely to observe it. If the “true” distribution of 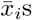 is described by some probability distribution function 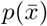, we expect to observe only the right part of the distribution, i.e.

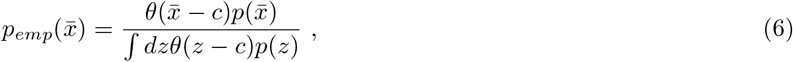

where *c* is the cutoff under which species are never observed because they are too rare (see also Supplementary Section S7). Note that, in reality *c* is not an hard cut-off. In this context, it refers to the minimal average abundance above which the error on the mean abundance due to sampling is negligible.

## Excluding competitive exclusion

A Gamma distributed AFD implies that all the species present in a community of a biome, are present in all the communities from that biome, and therefore, All the times a species is not observed is because of sampling errors. Since this result is very surprising, we tested it more carefully. It is important to underline that our claim is that competitive exclusion, at the taxonomic resolution at which species are defined in datasets we consider, is statistically insignificant (more rigorously defined below). We test this hypothesis in two independent ways.

The first way to test this hypothesis is to directly test its immediate prediction: if absence is a consequence of sampling, one should be able to predict occupancy of a species (the probability that a species is present) simply from its average and variance of abundance (together with the total number of reads of each sample). In particular, assuming a Gamma AFD, the occupancy of species *i* is given by

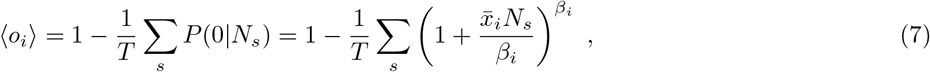

where *N*_*s*_ is the total number of reads in sample *s* (where *s* = 1, …, *T*) and 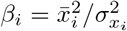. As shown in Figure 2 and in Supplementary Figure S3, this prediction well reproduces the observed occupancy across species. Note that the ability of a Gamma AFD to reproduce this pattern is also an indirect test of the hypothesis that the AFD is Gamma. For instance, Supplementary Figure S4 shows that assuming a Lognormal AFD would fail in reproducing the observed occupancy.

The second, more rigorous, way to test the hypothesis that (most) species are always present is to use model selection. In this context we want to compare two (or more) models that aim at describing the observed number of reads of each species starting from alternative hypothesis. In particular we compare a purely Gamma AFD with a zero inflated Gamma, which reads

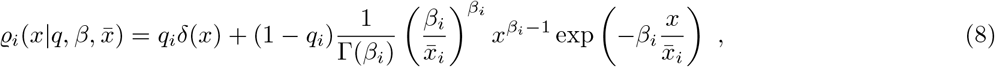

where *q*_*i*_ is the probability that a species is truly absent in a community and *δ*(·) is the Dirac delta distribution. Our goal is to test whether the *q*_*i*_s are significantly different from zero. Since the two models we are testing are nested, we compare the maximum likelihood estimator in the case *q*_*i*_ = 0 with the (maximum) likelihood marginalized over *q* (which has prior *μ*(*q*)). Given the number of reads 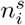 of species *i* in community *s* (with *N*_*s*_) total number of reads, we compute the ratio (see also Supplementary Section S4)

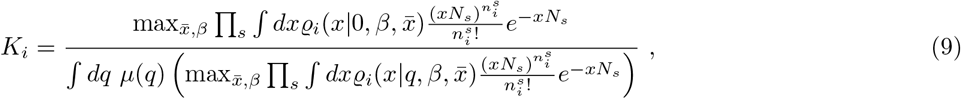

where *μ*(*q*) is a prior over *q*. If *K*_*i*_ > 1, the model with *q*_*i*_ = 0 is more strongly supported that the model with *q* ≠ 0. Under Beta prior with parameters 0.25 and 8 we obtained that *K*_*i*_ > 1 in 98.8% of the cases (averaged across datasets, ranging from 94.4% to 99.7%) and *K*_*i*_ > 100 in 97.5% cases (ranging from 92.8% to 99.2%). See Supplementary Section S4 for other choices of the prior and for a more detailed description of the results.

## Prediction of macroecological patterns

Given laws #1, #2, and #3, the probability to observe *n* reads of a randomly chosen species in a sample with *N* total reads is

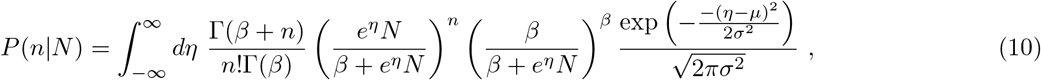

where 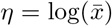. All the properties of species are fully specified by its mean abundance 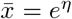. The probability of observing *k* reads of species with average abundance 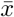 in a sample with *N* total number of reads is therefore

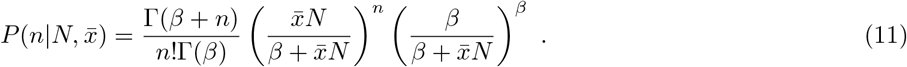

We now report the predictions for the patterns shown in Figure 3. For a full derivation of this and other patterns, see Supplementary Section S8.

The total number of observed species in a sample with *N* total number of reads can be easily calculated using equation 10. The probability of not observing an species is simply *P*(0|*N*). The expected number of distinct species ⟨*s*(*N*)⟩ in a sample with *N* reads is therefore

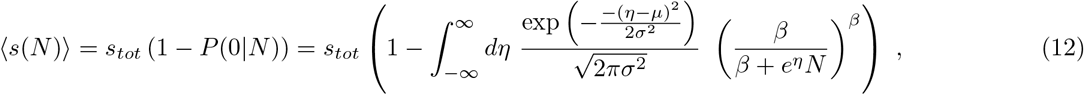

where *s*_*tot*_ is the total number of species in the biome (including unobserved ones, see Supplementary Section S7). Note that *s*_*tot*_ is (substantially) larger than *s_obs_*, the number of different species observed in the union of all the communities, which can instead be written as

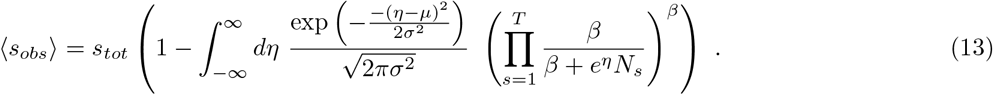

Figure 3a shows that the prediction of eq. 12 correctly matches the data (see also Supplementary Figure S9).

The Species Abundance Distribution (SAD), one of the most studied patterns in ecology and directly related to the Relative Species Abundance [22], is defined as the fraction of species with a given abundance. According to our model, the expected SAD is given by

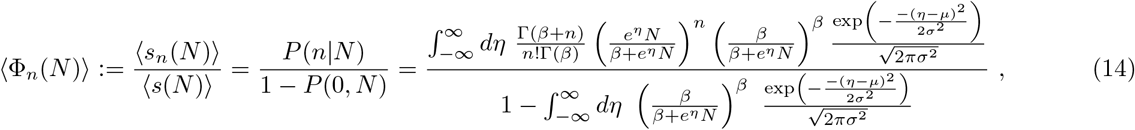

where ⟨*s*_*n*_(*N*)⟩ is the number of species with *n* reads in a sample with *N* total number of reads. The cumulative SAD is defined as

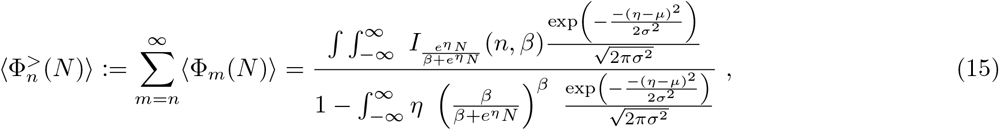

where *I*_*p*_(*n*, *β*) is the regularized incomplete Beta function. Figure 3b shows that the eq. S68 captures the empirical cumulative SAD (see also Supplementary Figure S22).

The occupancy probability is defined as the probability that a species is present in a given fraction of communities. This quantity has been extensively studied in a variety of contexts (from genomics [30] to Lego sets and texts [31]) and has been more recently considered in microbial ecology [24]. The three macroecological laws predict (see derivation in Supplementary Section S8)

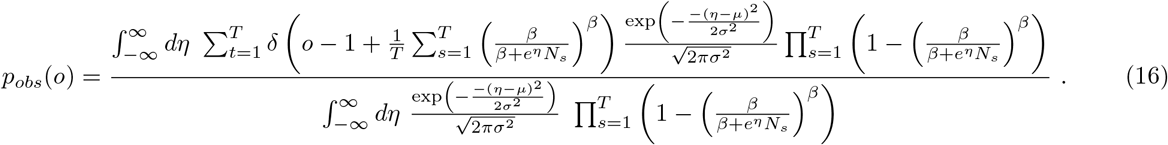

Figure 3b compares the prediction of eq. S61 with the data (see also Supplementary Figure S11).

Occupancy (the fraction of communities where a species is found) and abundance are not independent properties, and their relative dependence is often referred to as occupancy-abundance relationship [16] Given an average (relative) abundance 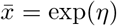, the expected occurrence is

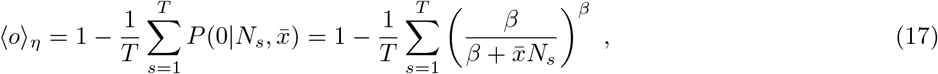

Figure 3d shows the comparison between data and predictions (see also Supplementary Figure S21).

## Transition Probabilities in Longitudinal Data

For longitudinal data, in addition to the stationary AFD, one can study the probability *ρ*_*i*_(*x*′, *t* + Δ*t*|*x*, *t*) that a species *i* has abundance *x*′ at time *t* + Δ*t*, given that the same species had abundance *x* at time *t*. Instead of focusing on the full distribution, we study its first two (conditional) central moments, i.e. the average and variance of the abundance at *t* + Δ*t* conditioned to abundance *x* at time *t*. In the analysis of the data we assume stationarity (the distribution *ρ*_*i*_(*x*′, *t* + Δ*t*|*x*, *t*) depends on Δ*t* but not on *t*). We test this assumption in section Supplementary Section S11.

We also assume that dynamics of different species are governed by similar equations that only differ in their parameters. We would like therefore to average over species, by properly rescaling their abundances. The average over species is potentially problematic, as it could add spurious effect to the conditional averages. For instance, only species with larger fluctuations would appear for extreme values of the initial abundance. In other to avoid these problems, instead of consider the actual abundance, we used its cumulative probability distribution value (calculated using the empirical AFD of each species), that we refer as “quantile abundance”. This is equivalent to rank the abundances of each species over communities and use the (relative) ranking of each community instead of the abundance. A value equal to 0 correspond to the lowest observed abundance, and a value equal to 1 to the highest. By definition, the quantile abundance is always uniformly distributed.

## Demographic stochasticity

Demographic stochasticity can reproduce a Gamma AFD. A birth, death and immigration process has a Gamma as stationary distribution [22]. In the limit of large populations sizes, it corresponds to the following equation [22]

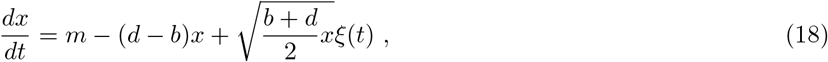

where *m* is the migration rate, while *b* and *d* are the per-capita birth and death rate. The Gaussian white noise term *ξ*(*t*) has mean zero and time-correlation ⟨*ξ*(*t*)*ξ*(*t*′)⟩ = *δ*(*t* − *t*′). The stationary distribution of this process turns out to be

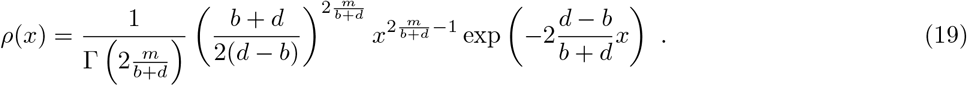

Mean and coefficient of variation of abundance are equal to 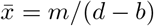 and 2*m*/(*b* + *d*), respectively.

More generally, we can assume that all the parameters are species dependent, and the population of species *i* is described by

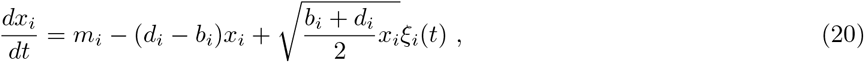

where we assume ⟨*ξ*_*i*_(*t*)*ξ*_*j*_(*t*′)⟩ = *δ*_*ij*_*δ*(*t* − *t*′).

Taylor’s Law and the Lognormal MAD together imply that *m*_*i*_/(*b*_*i*_ + *d*_*i*_) is constant while *m*_*i*_/(*d*_*i*_ − *b*_*i*_) varies on several orders of magnitude, corresponding to a non-trivial, and somewhat unnatural, relationship between migration rate and birth and death rate. If the number of individuals of the most abundance species is of the order of 10^9^ [30], the range of variability of (*b*_*i*_ + *d*_*i*_)/(*b*_*i*_ − *d*_*i*_) should be of the same order, implying a fine-tuned and unnatural scaling of parameters. It is important to underline, that the model of equation 20 can, in fact, for a proper parameterization, explain the observed variation of the data. But the choice of parameters explaining the empirical variation require for achieving this goal requires careful and unrealistic fine-tuning of the microscopic parameters.

## Stochastic Logistic Model

The Stochastic Logistic Model is defined as

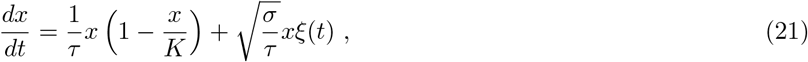

where *ξ*(*t*) is a Gaussian white noise term (mean zero and correlation ⟨*ξ*(*t*)*ξ*(*t*′)⟩ = *δ*(*t* − *t*′)), while the parameters 1/*τ*, *K* and *σ* are the intrinsic growth-rate, the carrying capacity and the coefficient of variation of the growth rate fluctuations. The stationary distribution of this process is

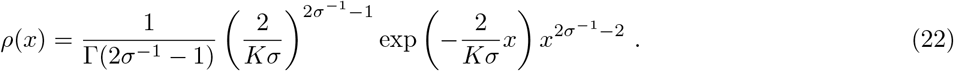

In general, we can assume that all the parameters are species dependent, and the population of species *i* is described by

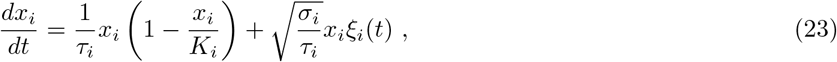

where we assume ⟨*ξ*_*i*_(*t*)*ξ*_*j*_(*t*′)⟩ = *δ*_*ij*_*δ*(*t*−*t*′). Taylor’s Law and the observed Lognormal MAD constraints the parameters value. Taylor’s Law requires *σ*_*i*_ = *σ* (independently of *i*), while the Lognormal MAD implies that the *K*_*i*_s are lognormally distributed.

The timescale *τ*_*i*_ does not affect stationary properties, but determines the timescale of relaxation to the stationary distribution. For small deviation of abundance from the average and for large times, the conditional expected abundance behaves as

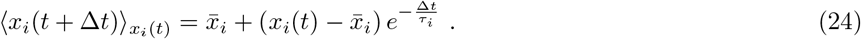

From the slopes of Figure 4g we can then determine the timescales *τ*_*i*_, which turn out to be approximately equal to 19 hours. In Figure 4 we assumed *τ*_*i*_ = 19 hours for all species.

