## Supplementary Material for "Laws of diversity and variation in microbial communities"

### Laws of diversity and variation in microbial communities — Supplementary Materials—

#### S1. DATA

All the datasets analyzed in this work were obtained from EBI Metagenomics [28] (now Magnify) and have been previously published. Raw data were processed under different version of EBI Metagenomics pipelines [28]. The consistency of results across studies and pipelines strongly support the generality of the conclusions. Supplementary Table S1 reports reference to the original works, Magnify pipeline and other information about each dataset. Note that the pipeline version 4.1 uses SILVA [32] to assign OTU classification, while previous versions of the pipeline use QIIME [33] and Greengenes [34]. In the following we will use species to refer to OTUs, defined accordingly to the methods referred above.

Supplementary Table S1: Description and references for the datasets used in this work.

| Biome ID | Type | EBI ID | Magnify ID | Pipeline Version | NCBI ID | Reference | # Samples T | Range Tot # Reads $N_s$ |
|---|---|---|---|---|---|---|---|---|
| glacier | c | ERP017997 | MGYS00001292 | 3.0 | PRJEB16145 | [35] | 30 | $[79765, 1104214]$ |
| gut1 | c | SRP056641 | MGYS00001056 | 2.0 | PRJNA275349 | [35] | 66 | $[13842, 102971]$ |
| gut2 | c | ERP015450 | MGYS00001556 | 3.0 | PRJEB13870 | [36] | 195 | $[24717, 614229]$ |
| lake | c | ERP012927 | MGYS00001669 | 3.0 | PRJEB11530 | [37] | 198 | $[57408, 350877]$ |
| oral1 | c | SRP056641 | MGYS00001056 | 2.0 | PRJNA275349 | [35] | 62 | $[10006, 138172]$ |
| river | c | ERP012927 | MGYS00001669 | 3.0 | PRJEB11530 | [37] | 188 | $[76042, 352675]$ |
| seawater | c | SRP128662 | MGYS00002437 | 4.1 | PRJNA429259 | [38] | 474 | $[11260, 492477]$ |
| sludge | c | ERP009143 | MGYS00001064 | 2.0 | PRJEB8105 | [37] | 575 | $[22255, 912713]$ |
| soil | c | SRP052295 | MGYS00000905 | 2.0 | PRJNA272333 | - | 112 | $[11352, 58219]$ |
| feces F4 | l | ERP021896 | MGYS00002184 | 4.1 | PRJEB19825 | [39] | 131 | $[21008, 51986]$ |
| feces M3 | l | ERP021896 | MGYS00002184 | 4.1 | PRJEB19825 | [39] | 334 | $[15047, 58463]$ |
| L_palm F4 | l | ERP021896 | MGYS00002184 | 4.1 | PRJEB19825 | [39] | 134 | $[12298, 34607]$ |
| L_palm M3 | l | ERP021896 | MGYS00002184 | 4.1 | PRJEB19825 | [39] | 365 | $[144, 48475]$ |
| R_palm F4 | l | ERP021896 | MGYS00002184 | 4.1 | PRJEB19825 | [39] | 134 | $[3214, 9052]$ |
| R_palm M3 | l | ERP021896 | MGYS00002184 | 4.1 | PRJEB19825 | [39] | 358 | $[135, 91953]$ |
| Tongue F4 | l | ERP021896 | MGYS00002184 | 4.1 | PRJEB19825 | [39] | 135 | $[5683, 12651]$ |

#### S2. SAMPLING

##### A. Notation

Let $N_s$ be the total number of reads in a biological sample $s$, and $n_i^s$ the number of reads belonging to species (or any other taxonomic classification) $i$. By definition $\sum_i n_i^s = N_s$ (if not all the reads are assigned to a species, we can introduce an unassigned category, such that $n_{\emptyset}^s$ is the number of unassigned reads).

We assume that the set of reads $\{n_\cdot^s\}$ is sample $s$ is produced by a (biased) sampling process. The probability $P_s(\{n.\}|N_s)$ of observing a given set of reads $\{n.\}$ conditioned to a total number of reads $N_s$ is multinomially distribute, i.e.,

$$P_s(\{n.\}|N_s) = \frac{N_s!}{\prod_i n_i!} \prod_i (p_i^s)^{n_i} \ , \tag{S1}$$

where $p_i^s$ is the actual frequency of species $i$ in sample $s$.

We are interested in the variability of abundance across samples. We can recapitualate this variability as fluctuations of the frequencies $p$s across samples, which are described by some (unknown) probability distribution $\rho(\{p.\})$. In this way, we can write equation S1 as

$$P(\{n.\}|N) = \int [dp] \ \rho(\{p.\}) \frac{N!}{\prod_i n_i!} \prod_i (p_i)^{n_i} \ . \tag{S2}$$

This equation disentangle two, important, sources of variability: the variability across samples/communities (described by $\rho(\{p.\})$) and the variability/noise due to sampling (decribed by the multinomial sampling).

Our goal is to access and study the properties of $\rho(\{p.\})$, i.e. we are interested in the fluctuations of the $p$s, how they are distributed and how they are correlated.

By marginalizing equation S2, without any additional assumption (e.g., independence), one obtains that the probability of observing $n$ reads of the OTU $i$ is

$$P_i(n|N) = \int dp \ \rho_i(p) \binom{N}{n} p^n (1-p)^{N-n} \ , \tag{S3}$$

where

$$\rho_i(p) = \int [dp] \quad \rho(\{p.\}) \delta(p_i - p) \ , \tag{S4}$$

where $\delta(\cdot)$ is the Dirac's delta function.

From equation S2, it is easy to obtain

$$\langle n_i \rangle_N := \sum_n P_i(n|N) n_i = \int [dp] \ \rho(\{p.\}) N p_i = N \langle p_i \rangle \ . \tag{S5}$$

Since the sampling effort (total number of reads) $N_s$ varies across samples, we cannot directly evaluate $\langle n_i \rangle_N$ from the data. On the other hand we can easily remove the effect of $N$, by computing $\langle n_i \rangle_N / N$. We can therefore easily estimate $\langle p_i \rangle$

$$\langle p_i \rangle \approx \frac{1}{T} \sum_{s=1}^{T} \frac{n_i^s}{N_s} \ . \tag{S6}$$

Applying the same concept to the second moment, we obtain

$$\langle n_i^2 \rangle_N = \int [dp] \ \rho(\{p.\}) (N p_i (1-p_i) + N^2 p_i^2) = N \langle p_i \rangle + N(N-1) \langle p_i^2 \rangle \ . \tag{S7}$$

We have therefore that

$$\frac{\langle n_i^2 \rangle_N - \langle n_i \rangle_N}{N(N-1)} = \langle p_i^2 \rangle \ , \tag{S8}$$

from which we can estimate

$$\langle p_i^2 \rangle \approx \frac{1}{T} \sum_s \frac{n_i^s(n_i^s - 1)}{N_s(N_s - 1)} \ . \tag{S9}$$

Note that, even if $N_s \gg 1$, it is in principle incorrect to evaluate $\langle p_i^2 \rangle$ as the sample mean of $(n_i^s/N_s)^2$. This approximation is justified only if all the $n_i^s \gg 1$.

##### B. Poisson approximation and compositional data

Sequencing data are compositional [29] and therefore their fluctuations are always relative: the equivalence $\sum_i p_i = 1$ constraints the fluctuations of species. If the abundance of a species goes up, the abundance of all the other species has to go down on average. This constraint is explicit in equation S1, or, equivalently in the probability of observing $n_i$ individuals of species $i$ given a relative abundance $p_i$

$$P(n_i|N, p_i) = \binom{N}{n_i} p_i^n (1 - p_i)^{N-n} \ . \tag{S10}$$

At this point, it is useful to consider that we are typically interested in the case where the number of reads is large, i.e. $N_s \gg 1$ (we considered only samples where $N_s > 10^4$, see table S1). Moreover, the most abundant species are a typically small fraction of the samples (i.e., $p_i \ll 1$). In this regime we can approximate the Binomial distribution which appears in equation S10 with a Poisson distribution, obtaining

$$P_i(n|N) = \int dx \ \rho_i(x) \frac{(xN)^n}{n!} e^{-xN} \ , \tag{S11}$$

which correspond to equation S3. The variable $x$ has the same interpretation as $p$: the relative abundance of a given species.

In principle the constraint $\sum_i x_i$ still holds. We want to show that the constraint can be relaxed to a milder condition, when it holds on average and $\sum_i \bar{x}_i = 1$. The joint distribution $\rho(\{x.\})$ can be written as

$$\rho(\{x.\}) = \frac{1}{Z} \tilde{\rho}(\{x.\}) \delta \left( \sum_{i=1}^{s} x_i - 1 \right) \ , \tag{S12}$$

where $s$ is the number of species and $\tilde{\rho}(\{x.\})$ is a distribution without the constraint $\sum_i x_i = 1$ and $Z$ is a normalization factor. The moment generating function is defined as $G(h.) = \int [dx] \ \rho(\{x.\}) \exp(i \sum_{j=1}^{s} h_j x_j)$. Using the integral representation of the delta function $\int dk \ \ \exp(ikx) = \delta(x)$, one obtains

$$G(\{h.\}) = \frac{1}{Z} \int [dx] \ \rho(\{x.\}) \exp(i \sum_{j=1}^{s} h_j x_j) = \frac{1}{Z} \int dk \ \ \int [dx] \ \tilde{\rho}(\{x.\}) e^{-ik + i \sum_{j=1}^{s} (h_j + k) x_j} = \frac{1}{Z} \int dk \ e^{-ik} \tilde{G}(\{h. + k\}) \ , \tag{S13}$$

where $\tilde{G}(\{h.\}) = \int [dx] \ \tilde{\rho}(\{x.\}) \exp(i \sum_{j=1}^{s} h_j x_j)$. If we expand $\tilde{G}(\{h. + k\})$ around $k = 0$, we obtain

$$\begin{aligned} G(\{h.\}) = & \frac{1}{Z} \int dk \ e^{-ik} \sum_{N=0}^{\infty} \sum_{\{n_1, n_2, \ldots, n_s\}} \delta_{\sum_i n_i, N} \frac{1}{N!} \frac{\partial^N G(\tilde{\{h.\}})}{\partial^{n_1} h_1 \partial^{n_2} h_2 \cdots \partial^{n_s} h_s} k^N = \\ = & \frac{1}{Z} \tilde{G}(\{h.\} \int dk \ e^{-ik} \sum_{N=0}^{\infty} \sum_{\{n_1, n_2, \ldots, n_s\}} \delta_{\sum_i n_i, N} \frac{1}{N!} \langle \tilde{x}_1^{n_1} \tilde{x}_2^{n_2} \ldots \tilde{x}_s^{n_s} \rangle (ik)^N = \\ = & \frac{1}{Z} \tilde{G}(\{h.\} \int dk \ e^{-ik} \tilde{G}_S(k) \ . \end{aligned} \tag{S14}$$

where $\tilde{G}_S(k)$ is equal to

$$\tilde{G}_S(k) = \int [dx] \; \tilde{\rho}(\{x.\}) \exp\left(ik \sum_i x_i\right) \quad . \tag{S15}$$

Using standard properties of the moment generating function, we have that

$$\tilde{G}_S(k) = \exp(ik \sum_{k=1}^{\infty} c_k) \quad , \tag{S16}$$

where $c_k$ is the $k$th cumulant of $\sum_i x_i$. For instance $c_1$ is the average of $\sum_i x_i$, $c_2$ is the variance and so on. These cumulants are calculated using the distribution $\tilde{\rho}(\{x.\})$ (which does not have constraints on the sum of the variables). If all the variables $x_i$ were independent, it is easy to show that the $k$-th cumulant of $\sum_{i=1}^{s} x_i$ scales with the total number of species $s$ as $c_k \sim s^{1-k}$. This scaling relationship is obtained assuming that all the cumulants of $sx_i$ are finite in the limit $s \to \infty$. In this case the leading term is therefore given by $k = 1$, and therefore, for large $s$ one obtains

$$\tilde{G}_S(k) \approx \exp(ikc_1) = \exp(ik \sum_{i=1}^{s} \bar{x}_i) \; , \tag{S17}$$

where

$$\bar{x}_i = \int [dx] \; \tilde{\rho}(\{x.\}) \; x_i \; , \tag{S18}$$

,

By inserting this expression in equation S14, we obtain

$$G(\{h.\}) \approx \frac{1}{Z} \tilde{G}(\{h.\} \int dk \; e^{-ik} \exp(ik \sum_{i=1}^{s} \bar{x}_i) = \frac{1}{Z} \tilde{G}(\{h.\} \delta \left(\sum_{i=1}^{s} \bar{x}_i - 1\right) \quad . \tag{S19}$$

Which translates into

$$\rho(\{x.\}) = \frac{1}{Z} \tilde{\rho}(\{x.\}) \delta \left(\sum_{i=1}^{s} \bar{x}_i - 1\right) \quad . \tag{S20}$$

which constrains the sum of the averages abundances and not on the sum of the random variables. In other words, if the number of species is large and abundances fluctuates independently, the constraints of equation S12 that impose that the sum of the random variables is equal to the unity can be relaxed to a constrain on the sum of the average abundances. If the abundances do not fluctuate dependently is not in principle true that $c_k \sim s^{1-k}$. The first cumulant (the average) is not affected by correlations, while the second cumulant is. In fact

$$c_2 = \langle \left(\sum_i x_i\right)^2 \rangle - \langle \left(\sum_i x_i\right) \rangle^2 = \sum_{ij} \langle x_i x_j \rangle - \sum_i \langle x_i \rangle \sum_j \langle x_j \rangle = \sum_i \sigma_{x_i}^2 + \sum_{i \neq j} \left(\langle x_i x_j \rangle - \langle x_i \rangle \langle x_j \rangle\right) \quad . \tag{S21}$$

Since the second cumulants of $sx_i$ is finite, then the variance $\sigma_{x_i}^2 \sim s^{-2}$ and therefore $\sum_i \sigma_{x_i}^2 \sim s^{-1}$. The second term contains a covariance and is therefore determined by correlation. We can always write the covariance as $\langle x_i x_j \rangle - \langle x_i \rangle \langle x_j \rangle = \rho_{ij} \sigma_{x_i} \sigma_{x_j}$. The product $\sigma_{x_i} \sigma_{x_j}$ scales as $s^{-2}$ and the sum over $i \neq j$ gives a contribution $\sim s^2$. In order to have $c_1$ dominating over $c_2$, we need therefore that the typical $\rho \sim s^{-\alpha}$ with $\alpha > 0$. In other words, if the correlations are weak enough (i.e. we do not have that every species is correlated with every other species), our approximation still holds for large enough number of species. For instance, if correlations are byproducts of interactions between species and a typical species only interacts with a finite number of species (which does not grow indefinitely as the number of species increases), $\rho \sim s^{-1}$ recovering a scaling $c_2 \sim s^{-1}$. Therefore, for large number of species, we expect correlations not to play a role in determining the constraint for species abundance fluctuations.

##### C. Moment generating function

In equation S7 we computed the second moment of the number of reads of a given OTUs, obtaining a non trivial dependence on the total number of reads $N_s$. Knowing this dependence allowed to remove the effect of sampling and to estimate the second moment of relative abundance from data obtained under different sampling efforts $N_s$ (eq. S9). It is straigthforward to generalize this calculation to the other moments. A more compact and efficient way to achieve the same goal is to estimate the moment generating function. We can calculate

$$\langle z^{n_i} \rangle_N = \sum_{n=0}^{\infty} z^{n_i} P(n_i|N) = \int dx_i \; \rho_i(x_i) \exp\left(N x_i (z-1)\right) = \langle \exp\left(N x_i (z-1)\right) \rangle \; . \tag{S22}$$

By introducing $q = N(z-1)$, we then obtain

$$\langle \left(1 + \frac{q}{N}\right)^{n_i} \rangle_N = \langle \exp\left(x_i q\right) \rangle \; . \tag{S23}$$

This suggests that we can remove the effect of the variability in the sampling effort and estimate the moment generating function of $\rho_i(x)$ as

$$G_i(q) = \langle \exp\left(x_i q\right) \rangle \approx \frac{1}{T} \sum_s \left(1 + \frac{q}{N_s}\right)^{n_i^s} \; , \tag{S24}$$

where $T$ is the total number of samples.

#### S3. LAW #1: FLUCTUATIONS OF OTUS ABUNDANCE ACROSS SAMPLES ARE GAMMA DISTRIBUTED

Figure S1 shows that the species that are present in all samples have Gamma distributed fluctuations of abundance, i.e.

$$\rho_i(x) = \frac{1}{\Gamma(\beta_i)} \left(\frac{\beta_i}{\bar{x}_i}\right)^{\beta_i} x^{\beta_i - 1} \exp\left(-\beta_i \frac{x}{\bar{x}_i}\right) \; . \tag{S25}$$

We chose to plot only the abundances of species always present because it is not obvious (a priori) how to treat abundances equal to zero. The instances where species are absent are in fact very likely due to sampling errors, which confound the shape of the AFD at low abundances. In section S4 we will see that this in fact happens in the vast majority of cases.

The average relative abundance $\bar{x}_i$ can be simply estimated using equation S6

$$\bar{x}_i \approx \frac{1}{T} \sum_{s=1}^{T} \frac{n_i^s}{N_s} \; . \tag{S26}$$

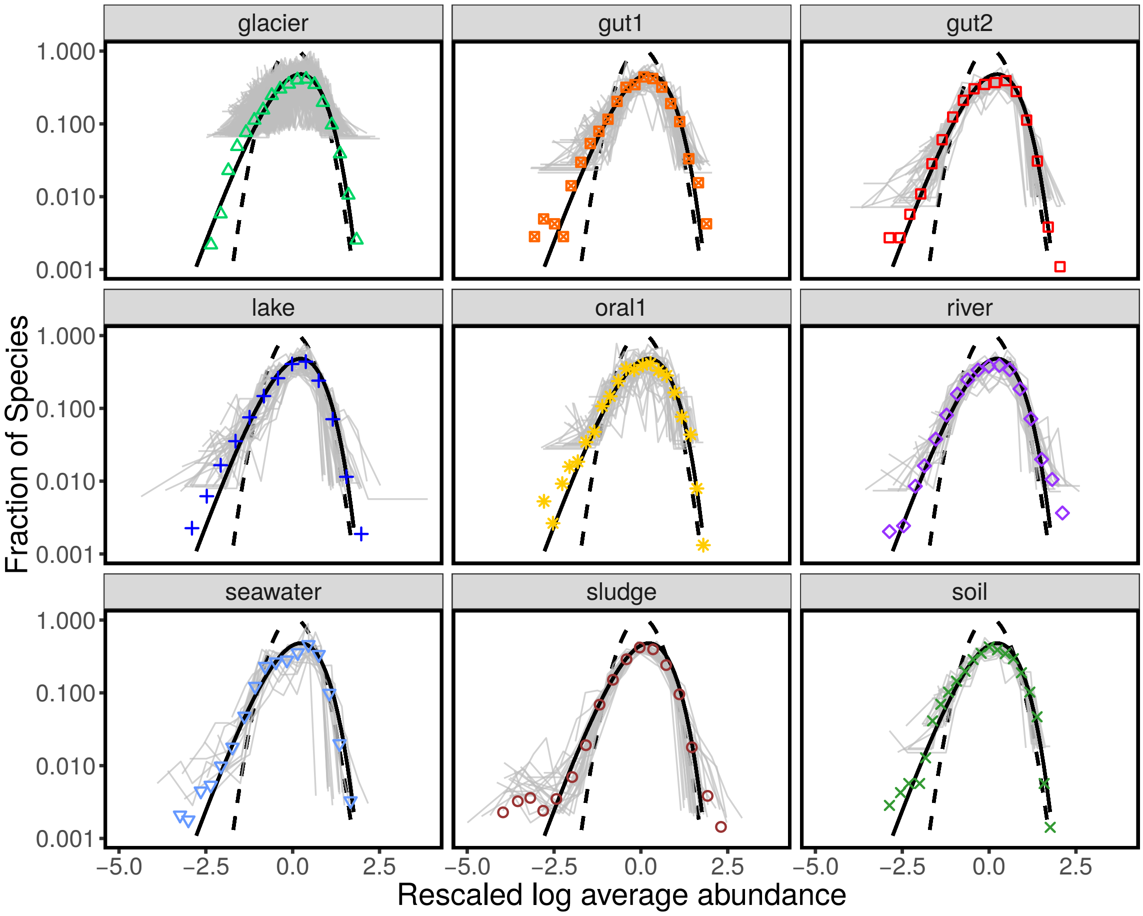


Supplementary Figure S1: **Fluctuations of species abundance.** These panels report exactly the same data shown in 1. For each biome, they were considered only the species present in all the communities. The logarithm of their relative abundances were rescaled (so to have mean zero and unitary variance). The panels report the distribution of these rescaled fluctuations for each biome. Colored points are calculated averaging over both communities and species (same as shown is figure 1). Gray lines are the distribution for individual species over communities. The black continuous line is a Gamma distribution and the black dashed line a Lognormal distribution.

Similarly, the parameter $\beta_i$ (which is related to the inverse of the coefficient of variation), can be obtained as

$$\beta_i \approx \left( \frac{1}{T} \sum_{s=1}^{T} \frac{n_i^s}{N_s} \right)^2 \left( \frac{1}{T} \sum_{s} \frac{n_i^s (n_i^s - 1)}{N_s (N_s - 1)} - \left( \frac{1}{T} \sum_{s=1}^{T} \frac{n_i^s}{N_s} \right)^2 \right)^{-1} . \tag{S27}$$

Since the variance of $x_i$ can be estimated as

$$\sigma_{x_i}^2 \approx \frac{1}{T} \sum_{s} \frac{n_i^s (n_i^s - 1)}{N_s (N_s - 1)} - \left( \frac{1}{T} \sum_{s=1}^{T} \frac{n_i^s}{N_s} \right)^2 , \tag{S28}$$

we can reqwrite $\beta_i$ as

$$\beta_i = \left( \frac{\bar{x}_i}{\sigma_{x_i}} \right)^2 . \tag{S29}$$

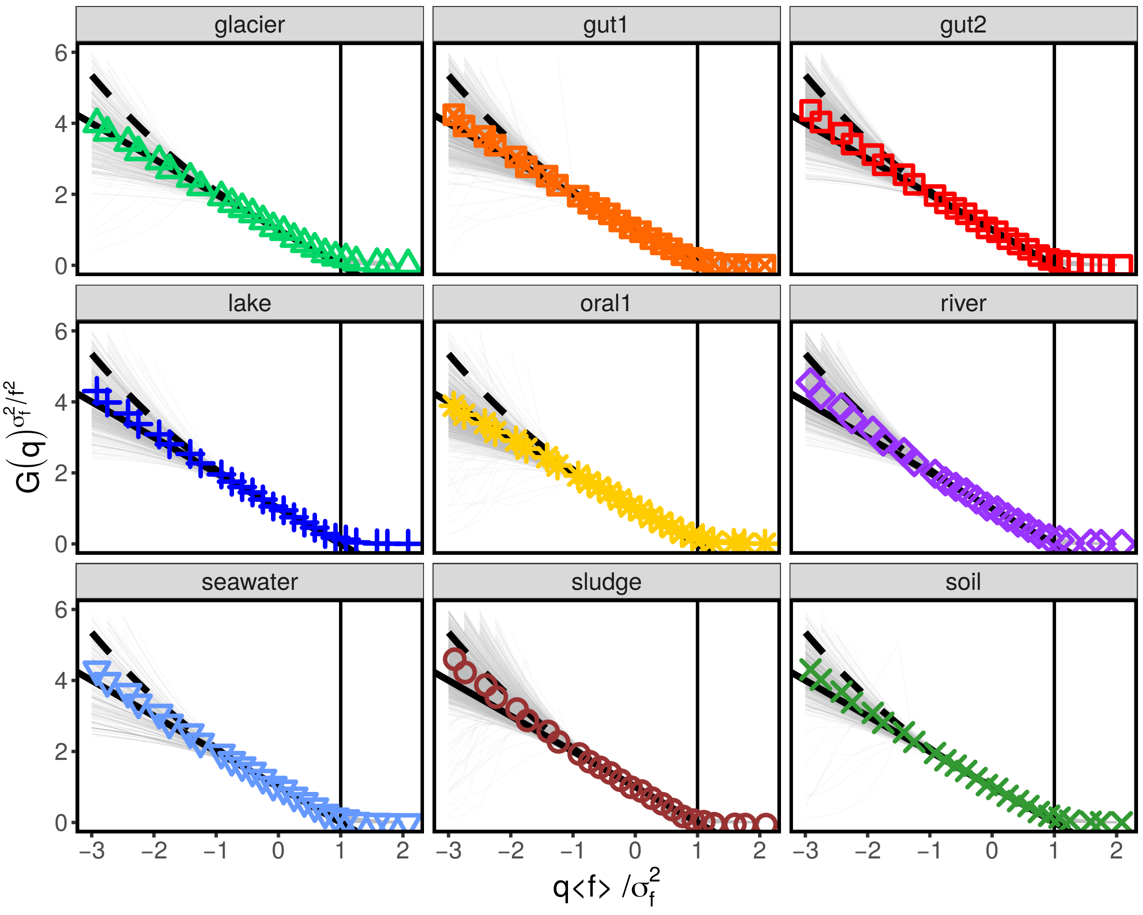


Supplementary Figure S2: **Moment generating function estimated from data.** The panels show the moment generating function estimated from the data using equation S32. The gray lines were obtained for individual species (with average average abundance $\bar{x}_i > 5 \cdot 10^{-5}$), while colored points are averages over species. The black solid line is the prediction for the Gamma distribution, while the dashed line is the one for a Lognormal.

The moment generating function of a Gamma distribution is

$$G_i(q) = \langle \exp(x_i q) \rangle = \left(1 - \frac{\bar{x}_i}{\beta_i} q\right)^{-\beta_i} , \tag{S30}$$

which can be estimated from data using equation S24. The moment generating function is species dependent, as is depends on $\bar{x}_i$ and $\beta_i$. On the other, it is easy to notice that $G_i(q)$ can be collapsed by rescaling both $q$ and $G_i$:

$$G_i\left(\beta_i \frac{q}{\bar{x}_i}\right)^{-\frac{1}{\beta_i}} = 1 - q , \tag{S31}$$

which is independent of $i$. We can therefore test whether $x_i$ is Gamma distributed, by checking that

$$\left(\frac{1}{S} \sum_s \left(1 + \frac{q}{N_s}\right)^{n_i^s}\right)^{-1/\beta_i} \approx (1 - q) . \tag{S32}$$

Figure S2 shows that the moment generating function estimated from the data using equation S32 is consistent with a Gamma AFD.

#### S4. EXCLUDING COMPETITIVE EXCLUSION

##### A. Prediction of occupancy from abundance average and variance

The fluctuations of abundance of species $i$ across samples are described by a Gamma distribution

$$\rho_i(x) = \frac{1}{\Gamma(r_i)\theta_i^{\beta_i}} x^{\beta_i - 1} \exp\left(-\beta_i \frac{x}{\bar{x}_i}\right) . \tag{S33}$$

The probability of observing $n_i$ reads of OTU $i$ in a sample with $N$ total number of reads is

$$P_i(n_i|N) = \frac{\Gamma(\beta_i + n_i)}{n_i!\Gamma(\beta_i)} \left(\frac{\bar{x}_i N}{\beta_i + \bar{x}_i N}\right)^{n_i} \left(\frac{\beta_i}{\beta_i + \bar{x}_i N}\right)^{\beta_i} , \tag{S34}$$

and, in particular, the probability of not observing species $i$ reads

$$P_i(0|N) = \left(1 + \frac{\bar{x}_i N}{\beta_i}\right)^{-\beta_i} . \tag{S35}$$

We define the occurrence of species $i$ as the fraction of samples where species $i$ is present, i.e.

$$o_i = \frac{1}{T}\sum_{s=1}^{T}\left(1 - \delta_{n_i^s,0}\right) = 1 - \frac{1}{T}\sum_{s=1}^{T}\delta_{n_i^s,0} , \tag{S36}$$

where the Kronecker delta $\delta_{k,0}$ is equal to 1 if $k = 0$ and zero otherwise. Using equation S35 we can calculate $\langle o_i \rangle$, the expected occurence of OTU $i$, which reads

$$\langle o_i \rangle = 1 - \frac{1}{T}\sum_{s=1}^{T} P_i(0|N_s) = 1 - \frac{1}{T}\sum_{s=1}^{T}\left(1 + \frac{\bar{x}_i N_s}{\beta_i}\right)^{-\beta_i} . \tag{S37}$$

Note that the two parameters $\bar{x}_i$ and $\beta_i$ are estimated independently of the occupancy, as they are function of the average relative abundance across samples and its variance only.

Figure S3 shows that a Gamma AFD, using which we obtained equation S37, correctly predicts empirical species' occupancies. One might wonder how sensitive is this success in reproducing the occupancy sensitive to the choice of a Gamma AFD. For instance, if the AFD was Lognormal (with parameters $m_i$ and $s_i$), we would expect

$$P_i(0|N_s) = \int d\eta \, \exp(-e^{\eta}) \frac{\exp\left(-\frac{(\eta - m_i)^2}{2s_i^2}\right)}{\sqrt{2\pi s_i^2}} , \tag{S38}$$

from which one can compute the expected occupancy $\langle o_i \rangle$ as $1 - \sum_{s=1}^{T} P_i(0|N_s)/T$. Figure S4 shows that a Lognormal AFD fails in reproducing the occupancies of species, always overestimating occupancies at intermediate values.

##### B. Model selection

In this section we compare a purely Gamma AFD with a zero inflated Gamma, which reads

$$\varrho_i(x|q, \beta, \bar{x}) = q_i\delta(x) + (1 - q_i)\frac{1}{\Gamma(\beta_i)}\left(\frac{\beta_i}{\bar{x}_i}\right)^{\beta_i} x^{\beta_i - 1} \exp\left(-\beta_i \frac{x}{\bar{x}_i}\right) , \tag{S39}$$

where $q_i$ is the probability that a species is truly absent in a community and $\delta(\cdot)$ is the Dirac delta distribution. The assumption behind this model is that if a species is present (which happens with probability $1 - q$) its abundance

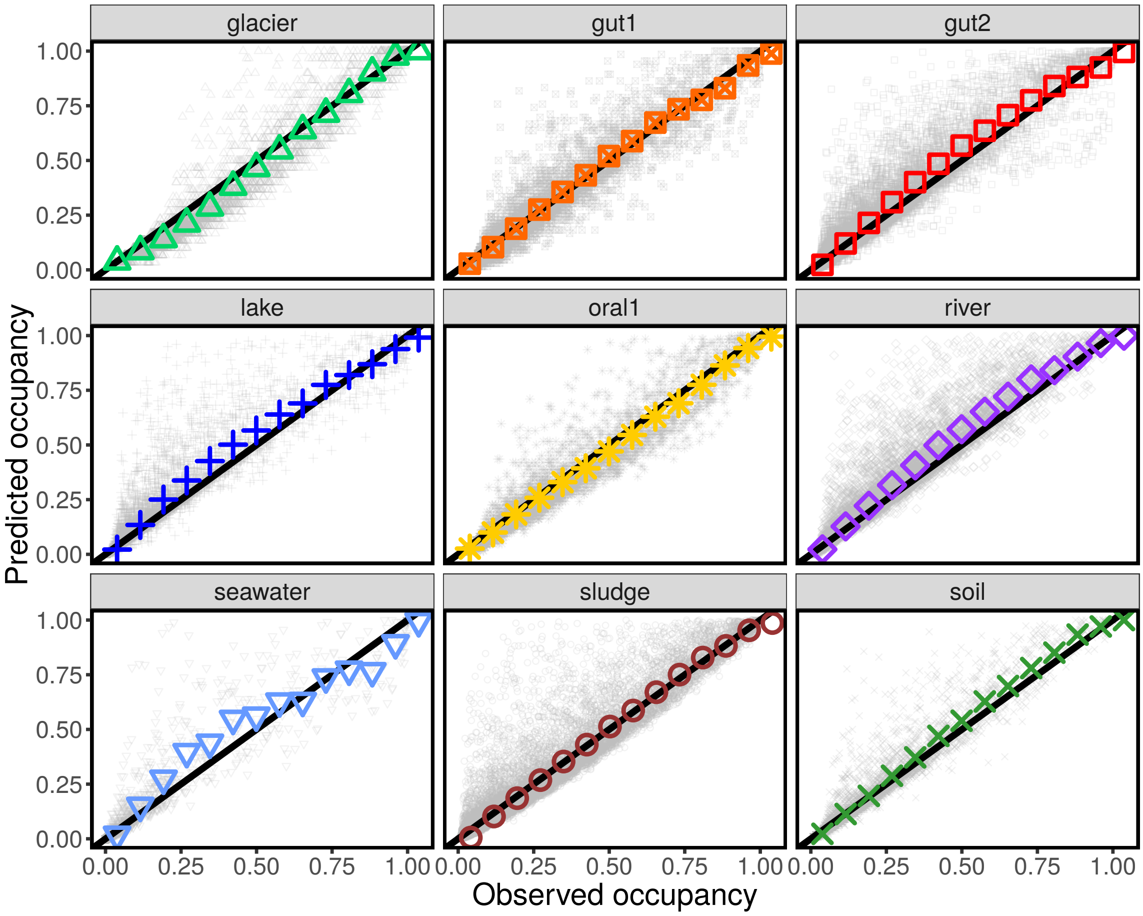


Supplementary Figure S3: **A Gamma AFD correctly predicts species' occupancies.** The occupancy is defined as the fraction of samples/commuties where a given species is found to be present. The predicted occupancy was obtained using equation S37, which assumes a Gamma AFD. Using the average and the variance of species' relative abundances, one can in fact estimate the parameters of the AFD and the probability that a species is not found in a sample/community, given the level of sampling. The black line is the 1 : 1 line, indicating a correct prediction. The gray points are individual species (no filter on average abundance was applied), while the colored points are averages over species.

fluctuations are Gamma distributed. Under this distribution the probability of observing $n_i$ reads for species $i$ in a sample with $N$ total number of reads is

$$P_i(n_i|N) = q_i \delta_{n_i,0} + (1-q_i) \frac{\Gamma(\beta_i + n_i)}{n_i! \Gamma(\beta_i)} \left( \frac{\bar{x}_i N}{\beta_i + \bar{x}_i N} \right)^{n_i} \left( \frac{\beta_i}{\beta_i + \bar{x}_i N} \right)^{\beta_i} , \tag{S40}$$

which reduces to equation S34 when $q_i = 0$ Note that $P_i(0|N_s)$ is always larger than $q_i$, as sampling errors are also present here and the probability of false negatives is always nonzero.

Our goal is to test whether the $q_i$s are significantly different from zero. Since the two models we are testing are nested, we introduce a prior $\mu(q)$ over the $q$ and we compare the maximum likelihood estimator in the case $q_i = 0$ with the (maximum) likelihood marginalized over $q$ with prior $\mu(q)$. Given the number of reads $n_i^s$ of species $i$ in

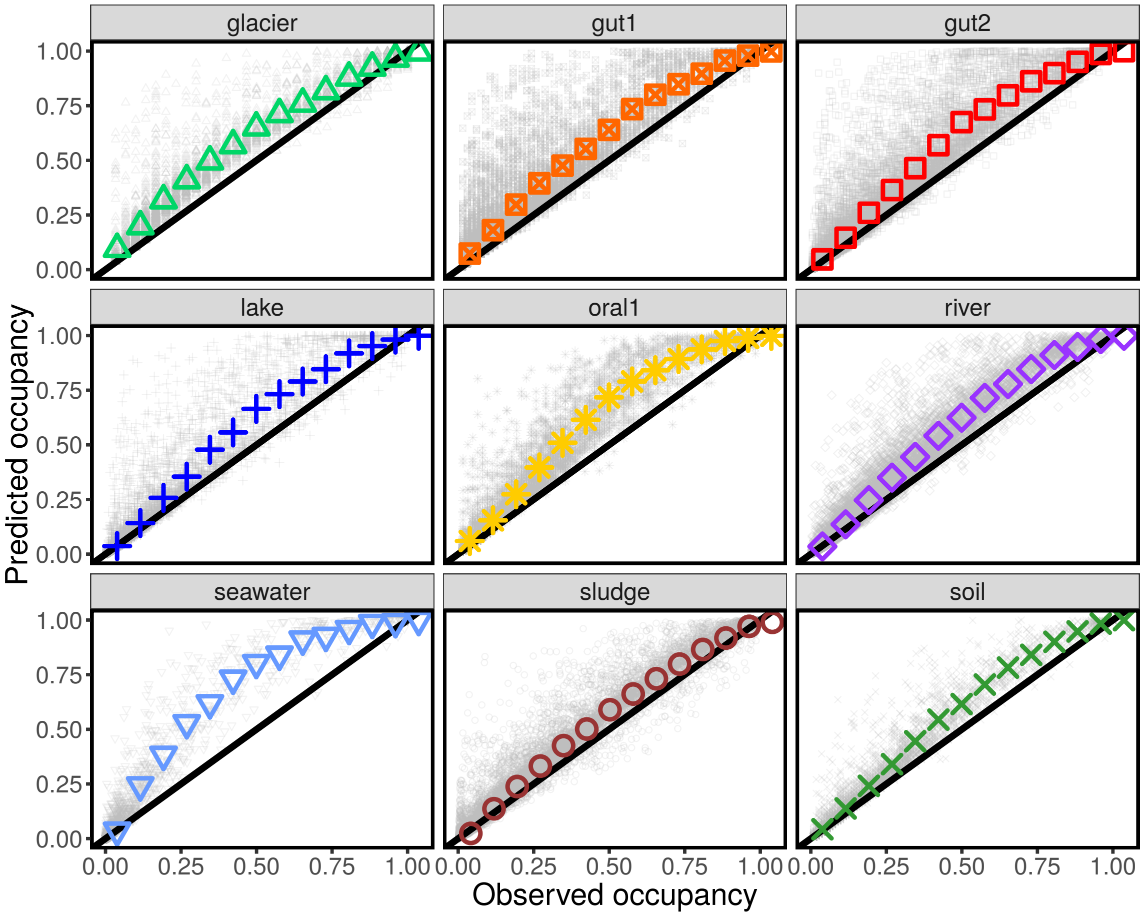


Supplementary Figure S4: **A Lognormal AFD fails in predicting species' occupancies.** The occupancy is defined as the fraction of samples/commuties where a given species is found to be present. The predicted occupancy was obtained using equation S38, which assumes a Lognormal AFD. Using the average and the variance of species' relative abundances, one can in fact estimate the parameters of the AFD and the probability that a species is not found in a sample/community, given the level of sampling. The black line is the 1 : 1 line, indicating a correct prediction. The gray points are individual species (no filter on average abundance was applied), while the colored points are averages over species.

sample $s$ (with $N_s$) total number of reads, we compute the ratio

$$K_i = \frac{\max_{\bar{x},\beta} \prod_s \int dx \varrho_i(x|0,\beta,\bar{x}) \frac{(xN_s)^{n_i^s}}{n_i^s!} e^{-xN_s}}{\int dq\ \mu(q) \left( \max_{\bar{x},\beta} \prod_s \int dx \varrho_i(x|q,\beta,\bar{x}) \frac{(xN_s)^{n_i^s}}{n_i^s!} e^{-xN_s} \right)} = \tag{S41}$$

$$= \frac{\max_{\bar{x},\beta} \prod_s \frac{\Gamma(\beta+n_i^s)}{n_i^s!\Gamma(\beta)} \left(\frac{\bar{x}N_s}{\beta+\bar{x}N_s}\right)^{n_i^s} \left(\frac{\beta}{\beta+\bar{x}N_s}\right)^{\beta}}{\int dq\ \mu(q) \left( \max_{\bar{x},\beta} \prod_s \left( q\delta_{n_i^s,0} + (1-q) \frac{\Gamma(\beta+n_i^s)}{n_i^s!\Gamma(\beta)} \left(\frac{\bar{x}N_s}{\beta+\bar{x}N_s}\right)^{n_i^s} \left(\frac{\beta}{\beta+\bar{x}N_s}\right)^{\beta} \right) \right)} .$$

If $K_i > 1$, the model with $q_i = 0$ is more strongly supported that the model with $q \neq 0$. We considered Beta prior

$$\mu(q) = \frac{\Gamma(a+b)}{\Gamma(a)\Gamma(b)} q^{a-1}(1-q)^{b-1} , \tag{S42}$$

which depends on two hyperparameters $a$ and $b$. In particular the average $q$ is equal to $a/(a+b)$.

For a given value of $q$ we numerically maximized $\prod_s \int dx \varrho_i(x|q,\beta,\bar{x}) \frac{(xN_s)^{n_i^s}}{n_i^s!} e^{-xN_s}$ over $\beta$ and $\bar{x}$. By calculating this

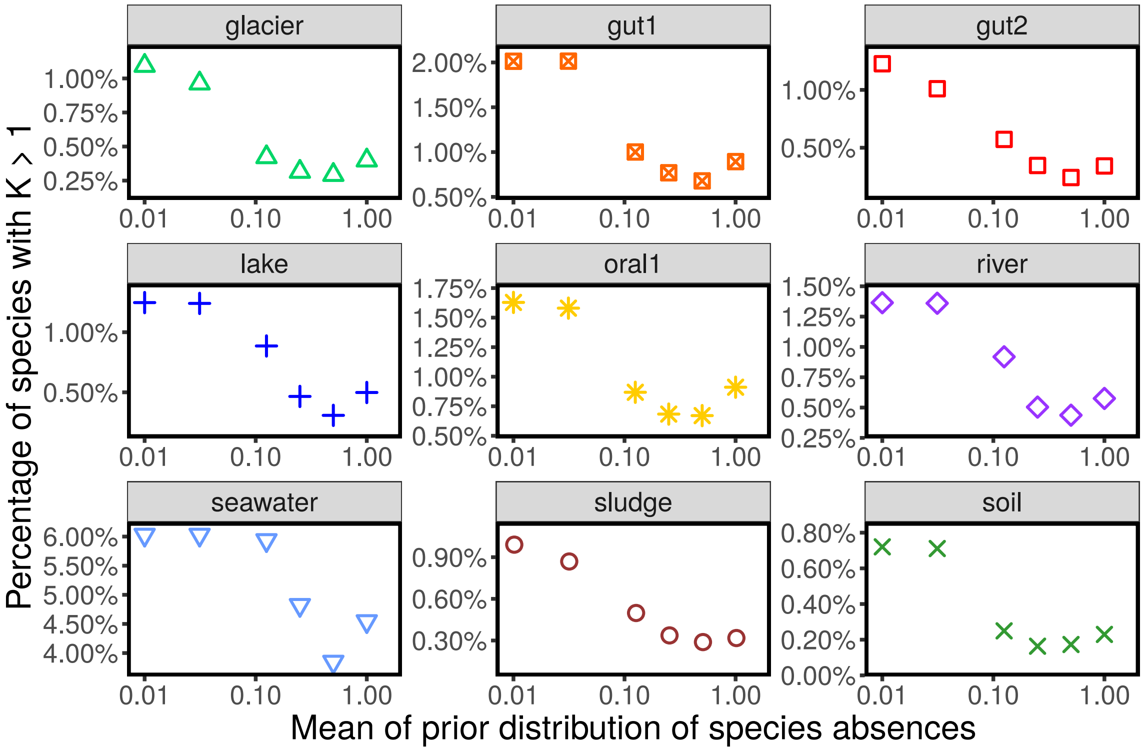


Supplementary Figure S5: **Fraction of species for which exclusion might be significant.** The plots show the fraction of species for which the inflated Gamma model (which allows true zeros in the abundance distribution) is more supported by the data then the standard Gamma model (which predicts that all the instances when a species is absent are due to sampling errors), measured as the fraction of species for which $K_i < 1$ (see definition in eq. S41). These values are plotted for different choices of the hyperparameters $a$ and $b$, showing that the fraction of species with $K_i < 1$ decreases with increasing the average probability of a true absence ($a/(a+b)$, on the x-axes).

maximum for $q = 0$ and comparing it with the averaged value of the maximum over Beta distributed $q$, we estimated $K_i$ for each species. Figure S5 reports the fraction of species for which $K_i < 1$, i.e., the fraction of species for which the inflated gamma is more statistically supported than the standard Gamma, suggesting a $q_i$ significantly different from zero. The value varies across biomes and over choices of $a$ and $b$, with tipical values of 1% to 10% of species displaying $K_i < 1$.

Interestingly the fraction of species with $K_i < 1$ decreases as the average $q_i$ (equal to $a/(a+b)$) increases. It is in fact important to notice that for $a \to 0$ (for $b > 0$), the Beta distribution tends to a Dirac delta distribution $\delta(q)$ and therefore the denominator and numerator of equation S41 become equal, implying $K = 1$. The fact that the fraction of species with $K_i < 1$ decreases monotonically with $a/(a+b)$ shows that the more the prior is concentrated around $q = 0$, the more likely the inflated Gamma distribution becomes, as it reduces to the standard Gamma distribution.

#### S5. LAW #2: TAYLOR'S LAW FOR ABUNDANCES FLUCTUATIONS

We showed that the fluctuations of OTU abundances across samples are well described by a Gamma distribution (what we called "Law #1"). The parameters of a Gamma distribution are fully specified by its mean and vari-

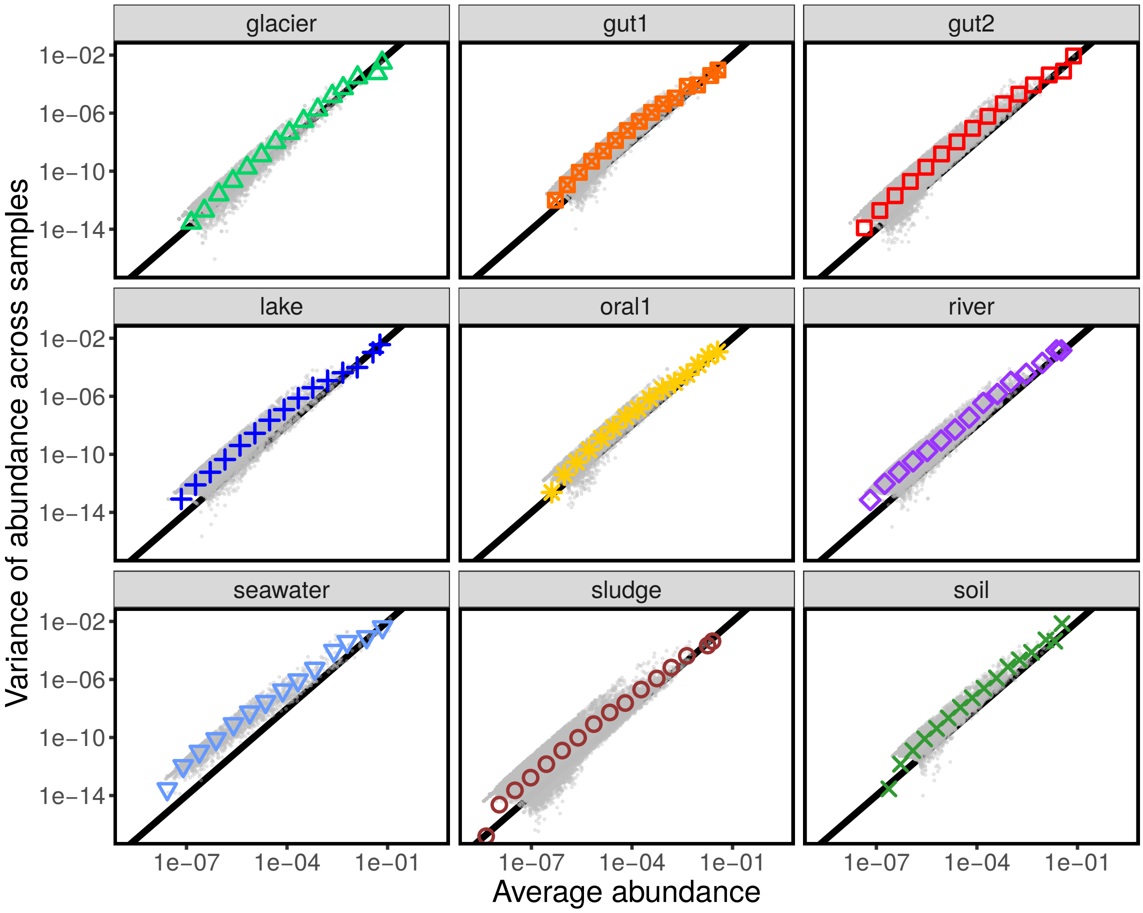


Supplementary Figure S6: **Taylor's law.** The panels show the relationship between mean and variance of abundance. The black line has slope 2, representing the quadratic relationship between variance and mean abundance.

ance. Knowing the mean and the variance of the relative abundance of a species, is therefore enough to specify the distribution of abundance and descending properties (e.g., the probability of being observed as shown in section S4 A).

In this section we explore the relation between mean and variance. Figure S6 shows that mean and variance are not independent across species. The variance is, in fact, proportional to the square of the mean. A relation of the type

$$\sigma_x^2 = \langle x \rangle^{2b} \,, \tag{S43}$$

is called Taylor's law and has been documented across multiple ecosystems [40]. As shown in Figure S6, we found $b = 1$, which implies that the coefficient of variation is constant. We can translate this observations into contraints on the parameters of the AFD. In fact, since $\beta_i$ depends only on the coefficient of variation, we can can neglect the fluctuations of $\beta_i$ across species.

#### S6. REPRODUCIBILITY OF AVERAGE ABUNDANCE

The fact that fluctuations of species abundance are well described by a Gamma AFD, reduce variation of each species' abundance to two parameters: mean and variance. Taylor's law, by establishing a link between mean and variance, implies that the average abundance of a species is the most important parameter to characterize its abundance fluctuations. In this section we show that the average summarize non-trivial biological information, being characteristic of a species in a set of similar environments. The average abundance could, in fact, be just a fitted value with little biological significance, not carrying any information about the environment where the species lived in. This could happen in two extreme way: the average is just a value independent on both species identity and biome (like it would be in neutral theory, see section S12) or the average depends on the species only (for instance, if the average abundance was a consequence of some technical choice in the experiment). Figure S7 shows the correlation of species' average abundances across biomes. For each pair of biomes we consider the species present in both biomes and we calculate the correlation between there average relative abundances. We observe both significant positive and negative correlations. Particularly high are the correlations between two different gut microbiome experiments and between river and lake microbiomes (both freshwater). This results indicates that average abundances are highly reproducible across experimental setups and have significant information about the particular set of environmental conditions.

#### S7. LAW #3: AVERAGE ABUNDANCES ARE LOGNORMALLY DISTRIBUTED

The fluctuations of species abundances across samples are fully specified by the average (relative) abundance. In this section we show that the average abundances are Lognormally distributed across species.

Since we are always dealing with a finite number of (finite) samples, not all the species are observed. We are interested in how the average abundances $\bar{x}_i$ are distributed across species. If a species is rare enough, (i.e., if $\bar{x}_i < c$, where $c$ is a cutoff) it becomes extremely unlikely to observe it. If the "true" distribution $\bar{x}_i$s is described by some probability distribution function $p(\bar{x})$, we expect to observe only the values larger than $c$

$$p_{emp}(\bar{x}) = \frac{\theta(\bar{x}-c)p(\bar{x})}{\int dz\theta(z-c)p(z)} \ , \tag{S44}$$

where $c$ is the cutoff under which species are never observed because they are too rare. Note that this cutoff is a probabilistic one (the probability of observing a species above this cutoff tends to one and is lower than one below).

Figure S8 shows that the distribution of $\bar{x}$ is consistent with a Lognormal distribution. In order to estimate the parameters of the Lognormal, we have to take into account the presence of the cutoff, as defined in equation S44. In other words, if $p(\bar{x})$ is Lognormal, the observed distribution of abundances will be

$$p_{emp}(\bar{x}) = \frac{\sqrt{2}}{\sqrt{\pi\sigma^2}\bar{x}}\theta(\bar{x}-c)\frac{\exp(-\frac{(\log(\bar{x})-\mu)^2}{2\sigma^2})}{\text{erfc}\left(\frac{\log(c)-\mu}{\sqrt{2}\sigma}\right)} \ . \tag{S45}$$

The parameters $\mu$ and $\sigma$ are unknown, and should be inferred from data, while $c$ is known (and depends on the number of samples and the sampling effort). The first and second moment of the log average abundances are given by

$$m_1 := \frac{1}{s_{obs}}\sum_i \log(\bar{x}_i) \ , \tag{S46}$$

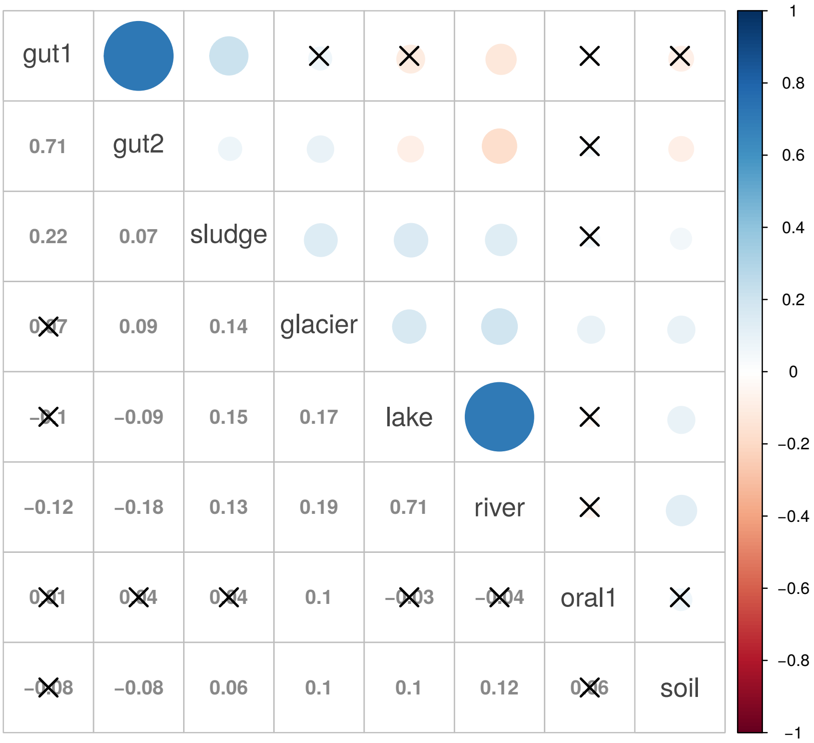


Supplementary Figure S7: **Correlation between average abundances across biomes.** The correlation plot shows the correlation between species average abundances across biomes. The colored circles represent the correlation value (also reported in the lower diagonal part). Crossed circles/values are non-significant correlations (the ones with p-value larger than 0.001). Note that the sample size varies across pairs of biomes as one can consider only species which appeared in both biomes. Seawater was excluded from this plot as the OTU picking method does not match with the ones of other biomes.

and

$$m_2 := \frac{1}{s_{obs}} \sum_i \left(\log(\bar{x}_i)\right)^2 \ , \tag{S47}$$

where $s_{obs}$ is the total number of OTUs observed across samples (with log average abundance larger than the cutoff $c$). It turns out that the maximum likelihood estimate of $\mu$ and $\sigma$ is the solution of the following system of equations

$$\begin{cases} m_1 = \dfrac{\sqrt{\frac{2}{\pi}}\sigma e^{-\frac{(\log(c)-\mu)^2}{2\sigma^2}}}{\mathrm{erfc}\left(\frac{\log(c)-\mu}{\sqrt{2}\sigma}\right)} + \mu \\ m_2 - m_1^2 = \sigma^2 - (m_1 - \mu)(\log(c) - m_1) \end{cases} \tag{S48}$$

The species below an average log abundance $c$ were not observed. The total number of species $s_{tot}$ can be inferred calculating the probability of observing a species, by assuming that the abundances of non observed species is also lognormally distributed, i.e., to be more precise, that the true distribution of average abundances is

$$p_{true}(\bar{x}) = \frac{\sqrt{2}}{\sqrt{\pi\sigma^2}\bar{x}} \exp\left(-\frac{(\log(\bar{x})-\mu)^2}{2\sigma^2}\right) \ . \tag{S49}$$

In this case the expected number of species $\langle s_{obs}\rangle$ with log abundance larger than $c$ is given by

$$\langle s_{obs}\rangle = s_{tot}\int_c^\infty d\bar{x}\ p_{true}(\bar{x}) = s_{tot}\,\mathrm{erfc}\left(\frac{\log(c)-\mu}{\sqrt{2}\sigma}\right) \ . \tag{S50}$$

We inferred the total number of species $s_{tot}$ by inverting this equation to obtain

$$s_{tot} = \frac{\langle s_{obs}\rangle}{\mathrm{erfc}\left(\frac{\log(c)-\mu}{\sqrt{2}\sigma}\right)} \ , \tag{S51}$$

and by using the empirical value of $s_{obs}$. The inferred values of $\mu$, $\sigma$ and $s_{tot}$ (together with the value of $\beta$) are reported in table S2.

Note that the true values of $\bar{x}_i$ are constrained by $\sum_{i=1}^{s_{tot}} \bar{x}_i = 1$, which implies a contraint on the value of parameters. In fact, we have that

$$\frac{1}{s_{tot}} = \frac{1}{s_{tot}}\sum_{i=1}^{s_{tot}} \bar{x}_i = \int_0^\infty d\bar{x}\ p_{true}(\bar{x})\ \bar{x} = \exp\left(\mu + \frac{\sigma^2}{2}\right) \ , \tag{S52}$$

which translated in the constraint

$$s_{tot}\exp\left(\mu + \frac{\sigma^2}{2}\right) = 1 \ . \tag{S53}$$

We did not impose this contraint in the inference of the parameters, as it leads to very unstable result. For instance, one could use equation S53 to infer $s_{tot}$ from $\mu$ and $\sigma$. A relatively small error in $\mu$ and/or $\sigma$ would be strongly amplified in the estimate of $s_{tot}$. We checked numerically (by generating samples from a constained lognormal distribution) that estimating independently the three parameters leads to more accurate results in the estimate of $\mu$, $\sigma$ and $s_{tot}$ at the expenses of imposing the constraint of equation S53 exactly.

#### S8. MACROECOLOGICAL PATTERNS ARE PREDICTED BY LAWS #1, #2 AND #3

Given laws #1, #2, and #3, the probability to observe $n$ reads of a randomly chosen OTUs in a sample with $N$ total reads is

$$P(n|N) = \int d\eta\ \frac{\Gamma(\beta+n)}{n!\Gamma(\beta)}\left(\frac{e^\eta N}{\beta+e^\eta N}\right)^n\left(\frac{\beta}{\beta+e^\eta N}\right)^\beta \frac{\exp\left(-\frac{-(\eta-\mu)^2}{2\sigma^2}\right)}{\sqrt{2\pi\sigma^2}} \ , \tag{S54}$$

where $\eta = \log(\bar{x})$ and where we are calculating the distribution over all the $s_{tot}$ species (i.e., including the unobserved species).

All the properties of a species are fully specified by its mean abundance $\bar{x} = e^\eta$. The probability of observing $k$ reads of a species with log average abundance $\eta$ in a sample with $N$ total number of reads is therefore

$$P(n|N,\eta) = \frac{\Gamma(\beta+n)}{n!\Gamma(\beta)}\left(\frac{e^\eta N}{\beta+e^\eta N}\right)^n\left(\frac{\beta}{\beta+e^\eta N}\right)^\beta \ . \tag{S55}$$

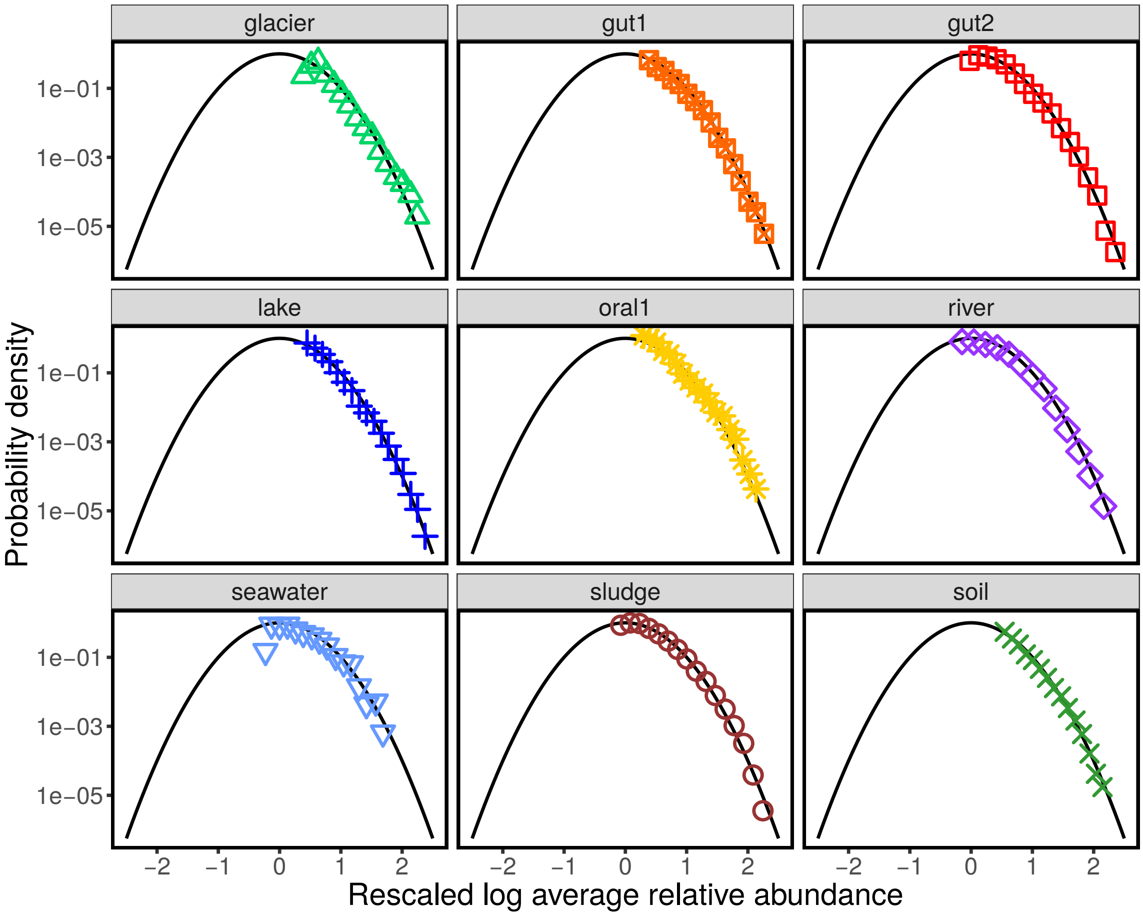


Supplementary Figure S8: **The Mean Abundance Distribution (MAD) is Lognormal.** The panels show the distribution of the logarithm of the average abundance $\log \bar{x}_i$. A Lognormal MAD corresponds to normally distributed $\bar{x}_i$. The black line is standardized gaussian distribution (mean zero and variance one). The log average abundances were rescaled as $z_i = (\log \bar{x}_i - \mu)/\sigma$, where $\mu$ and $\sigma$ were obtained for each biome from equation S48. The panels show that the rescaled log average abundances $z_i$ are distributed according to a standard normal, implying that the average abundances $\bar{x}_i$ are lognormally distributed.

##### A. Number of observed species vs total number of reads

The total number of observed species in a sample with $N$ total number of reads can be easily calculated using equation S54. The probability of not observing a species is simply $P(0|N)$. The expected number of distinct OTUs $\langle s(N) \rangle$ in a sample with $N$ reads is therefore

$$\langle s(N) \rangle = s_{tot} \left(1 - P(0|N)\right) = s_{tot} \left( 1 - \int d\eta \, \frac{\exp\left(-\frac{-(\eta-\mu)^2}{2\sigma^2}\right)}{\sqrt{2\pi\sigma^2}} \left( \frac{\beta}{\beta + e^{\eta} N} \right)^{\beta} \right) . \tag{S56}$$

Figure S9 compares the empirical relation between number of species and total number of reads with the prediction of equation S57.

Supplementary Table S2: Estimate of parameters across biomes and datasets.

| Biome ID | $s_{tot}$ | $\mu$ | $\sigma$ | $\beta$ |
|---|---|---|---|---|
| glacier | 20221 | -18.5 | 4.1 | 1.3 |
| gut1 | 35469 | -16.1 | 3.5 | 0.4 |
| gut2 | 50186 | -17.2 | 3.8 | 0.3 |
| lake | 46912 | -19.8 | 4.4 | 0.4 |
| oral1 | 29149 | -17.3 | 4.1 | 0.4 |
| river | 20833 | -14.2 | 2.7 | 0.3 |
| seawater | 3336 | -16.2 | 4.6 | 0.2 |
| sludge | 56671 | -17.5 | 3.7 | 0.6 |
| soil | 29387 | -17.1 | 3.8 | 0.38 |
| feces F4 | 1742 | -22.1 | 7.0 | 3.2 |
| feces M3 | 5575 | -28.0 | 8.5 | 1.2 |
| L_palm F4 | 3429 | -14.7 | 4.1 | 0.7 |
| L_palm M3 | 3179 | -14.8 | 4.0 | 0.3 |
| R_palm F4 | 2743 | -13.7 | 3.8 | 0.9 |
| R_palm M3 | 6547 | -18.4 | 5.2 | 0.3 |
| Tongue F4 | 1708 | -22.6 | 7.3 | 1.5 |
| Tongue M3 | 2368 | -23.9 | 7.4 | 1.4 |

##### B. Shannon index

Given a sample $s$ with $N_s$ total number of reads and with $n_i^s$ reads of OTU $i$, the Shannon diversity index is defined as

$$H_s = -\sum_{i \in s} \frac{n_i^s}{N_s} \log\left(\frac{n_i^s}{N_s}\right) . \tag{S57}$$

The expected Shannon index is therefore given by

$$\begin{aligned}\langle H(N)\rangle &= -s_{tot} \sum_{n>0} \frac{n}{N} \log\left(\frac{n}{N}\right) P(k|N) = \\ &= -s_{tot} \int d\eta \sum_{n>0} \frac{n}{N} \log\left(\frac{n}{N}\right) \frac{\Gamma(\beta+n)}{n!\Gamma(\beta)} \left(\frac{e^\eta N}{\beta + e^\eta N}\right)^n \left(\frac{\beta}{\beta + e^\eta N}\right)^\beta \frac{\exp\left(-\frac{-(\eta-\mu)^2}{2\sigma^2}\right)}{\sqrt{2\pi\sigma^2}} .\end{aligned} \tag{S58}$$

Figure S10 compares the predictions of equation S58 with data.

##### C. Occupancy distribution

The occupancy of a species is the fraction of samples where that species is present. The occupancy distribution is the probability distribution of observing a species with a given occupancy. The probability of observing a species with log average abundance $\eta$ in a sample with $N$ reads is $1 - P(0|N, \eta)$, where $P(n|N, \eta)$ is defined in equation S55. Let

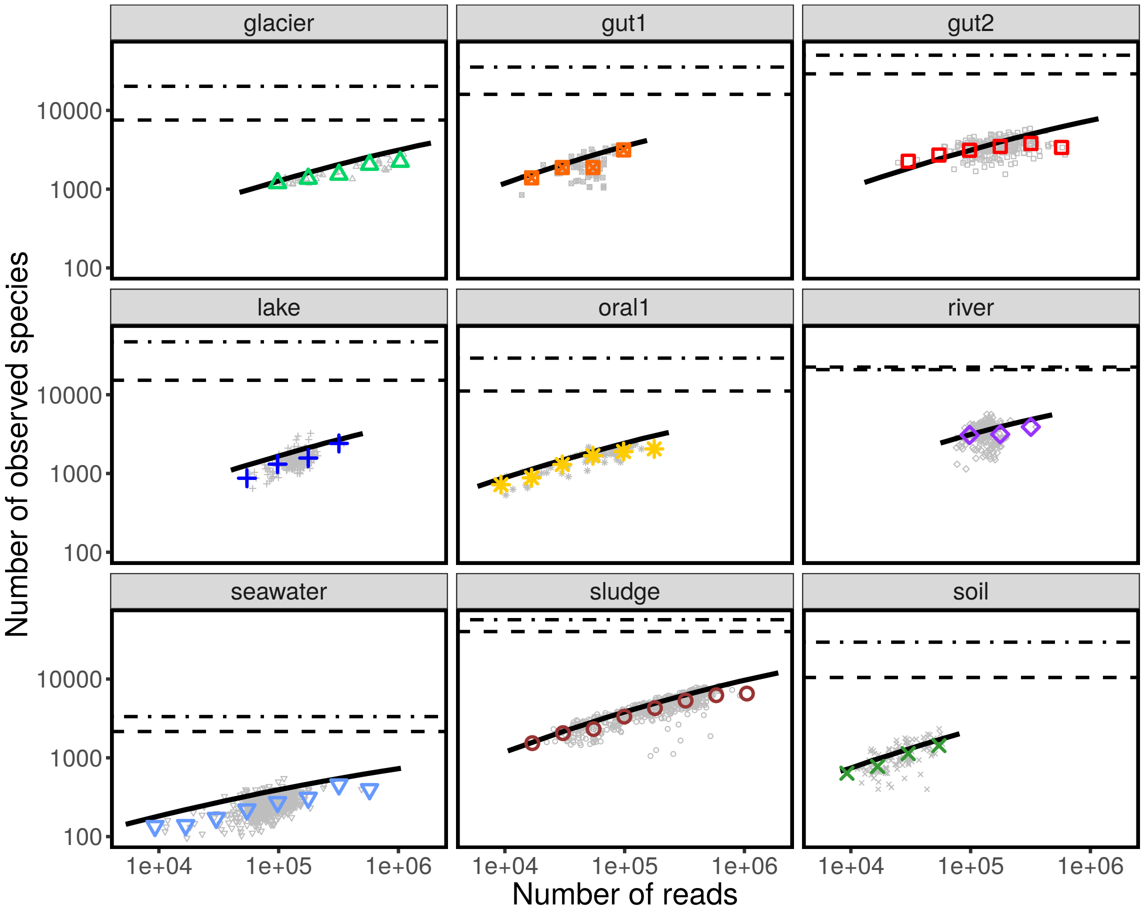


Supplementary Figure S9: **Number of observed species vs. total number of reads.** The number of observed species in a sample/community, depends on the total number of sequences sampled $N_s$. The more sequences are samples, the more likely it is to find new species. The gray points report the number of species observed in each community, while the colored symbols are averaged over communities in a range of $N_s$. The black dashed line represent $s_{obs}$, the total number of observed species across all the communities in that biome. The black dot-dashed line reports the value of the inferred value of $s_{tot}$, which includes also the species that have not been observed in that biome. The solid black line is the prediction, reported in equation S57, obtained by combining the three macroecological laws. Equation S57 correctly predicts the typical values of observed number of species, as well as the quantitative relationship between species and number of sequences.

us define $\chi_s(\eta)$, which is equal to 1 if an OTU with log average abundance $\eta$ is present in sample $s$ (which happens with probability $1 - P(0|N_s, \eta)$) and zero otherwise. The probability $p(o|\eta)$ that a species has occupancy $o$ is given by

$$p(o|\eta) = \sum_{\{\chi_1, \chi_2, \ldots, \chi_s, \ldots, \chi_T\}} \left[ \delta\left( o - \frac{1}{T} \sum_s \chi_s \right) \prod_s \left( \delta_{\chi_s,1} \left(1 - P(0|N_s, \eta)\right) + \delta_{\chi_s,0} P(0|N_s, \eta) \right) \right] . \tag{S59}$$

The number of samples $oT$ where a given OTUs is present is therefore given by a Poisson Binomial distribution, which describes the distribution of the sum of independent non-identically distributed Bernoully random variables. Using

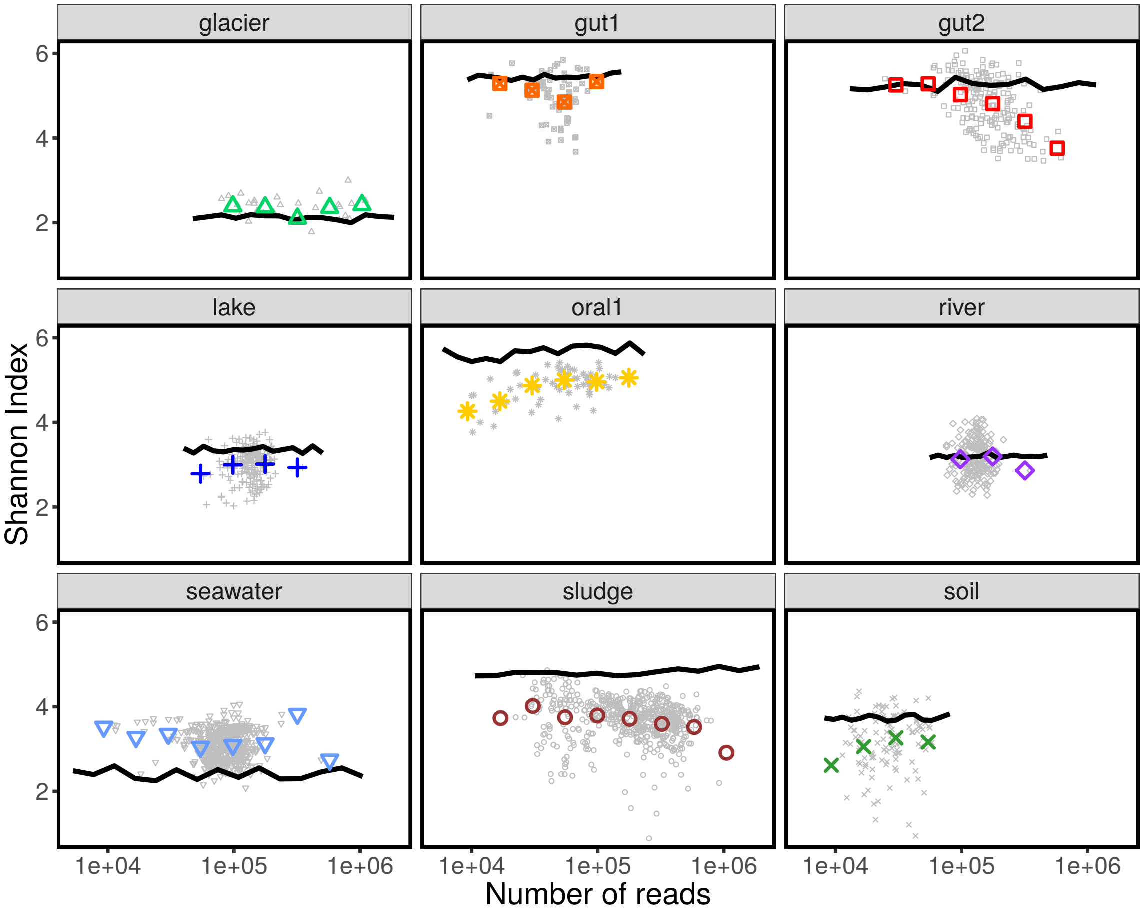


Supplementary Figure S10: **Shannon index vs. total number of reads.** The Shannon index is a measure of diversity which explicitely takes into account the distribution of abundances of species. The Shannon index is defined as the (Shannon) entropy of the probability that a random individual belongs to a given species. The gray points report the number of species observed in each community, while the colored symbols are averaged over communities in a range of number of reads $N_s$. The solid black line is the prediction, reported in equation S58, obtained by combining the three macroecological laws.

the Fourier transform of the Poisson Binomial distribution, we can write

$$p(o|\eta) = \sum_{t=0}^{T} \delta(o - t/T) \frac{1}{T+1} \sum_{l=0}^{T} e^{-\frac{2\pi i}{T+1} lt} \prod_{s=1}^{T} \left( P(0|N_s, \eta) + e^{\frac{2\pi i}{T+1} l} \left(1 - P(0|N_s, \eta)\right) \right) . \tag{S60}$$

The distribution of occupancy across species $p_{obs}(o)$ can be obtained by averaging S60 over $\eta$. Since we can observe only the OTUs with $o > 0$, we havce to restrict the summation in eq. S60 only over $t > 0$ and to normalize the distribution over the probability of observing a given number of species. The result of this calculation reads

$$p_{obs}(o) = \frac{\int d\eta \, \sum_{t=1}^{T} \delta(o - t/T) \frac{1}{T+1} \sum_{l=0}^{T} e^{-\frac{2\pi i}{T+1} lt} \prod_{s=1}^{T} \left( \left( \frac{\beta}{\beta + e^{\eta} N_s} \right)^{\beta} + e^{\frac{2\pi i}{T+1} l} \left( 1 - \left( \frac{\beta}{\beta + e^{\eta} N_s} \right)^{\beta} \right) \right) \frac{\exp\left( - \frac{-(\eta-\mu)^2}{2\sigma^2} \right)}{\sqrt{2\pi\sigma^2}}}{\int d\eta \, \frac{\exp\left( - \frac{-(\eta-\mu)^2}{2\sigma^2} \right)}{\sqrt{2\pi\sigma^2}} \, \prod_{s=1}^{T} \left( 1 - \left( \frac{\beta}{\beta + e^{\eta} N_s} \right)^{\beta} \right)} . \tag{S61}$$

This equation can be simplified by assuming that the occupancy of each species is equal to its mean value, which

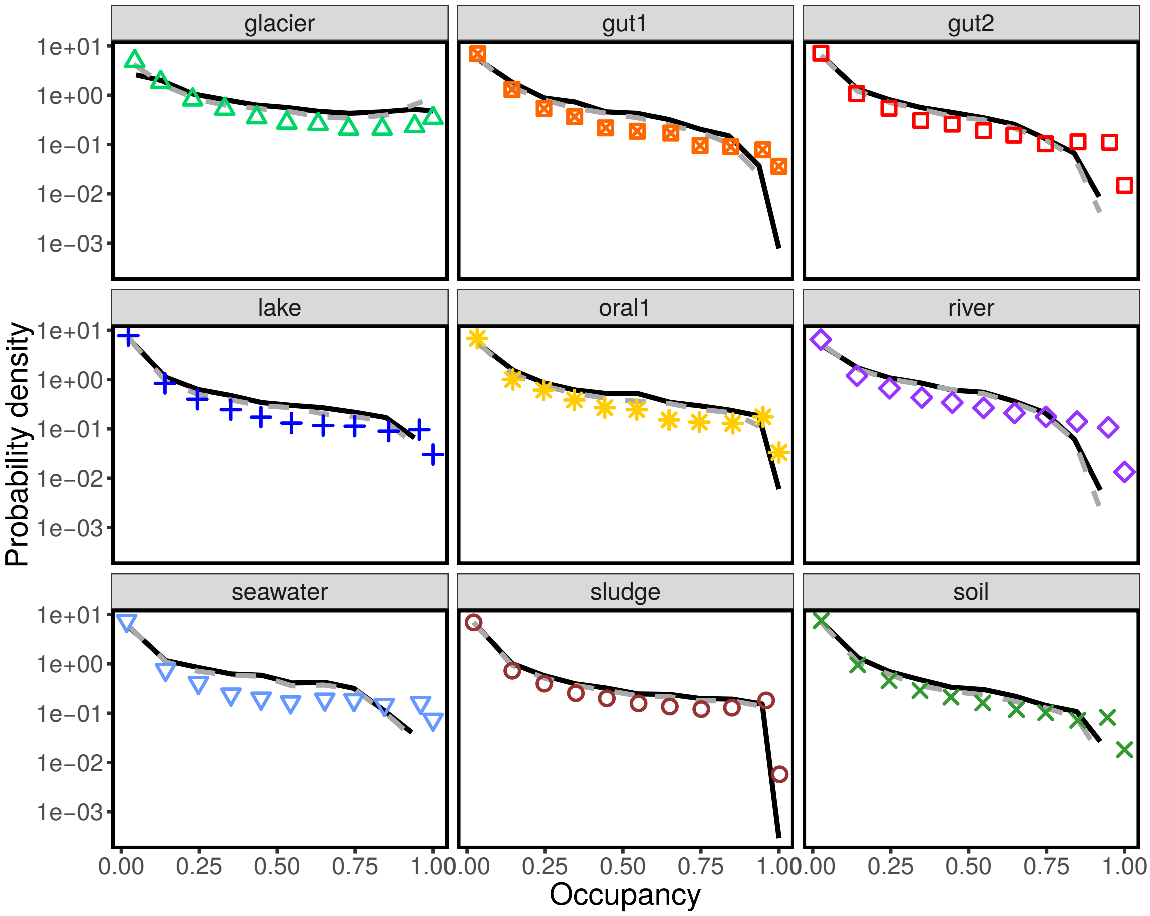


Supplementary Figure S11: **Distribution of occupancy.** The colored symbols report the distribution of species' occupancy in each biome. The solid black line is the prediction, reported equation S61, obtained by combining the three macroecological laws. The gray dashed line is the approximation to the prediction reported in equation S63.

is justified if the number of samples is large. This assumption corresponds to write

$$p(o|\eta) = \delta(o - \langle t \rangle / T) = \delta(o - \langle o \rangle_\eta) = \delta \left( o - (1 - \frac{1}{T} \sum_{s=1}^{T} P(0|N_s, \eta)) \right) . \tag{S62}$$

By averaging over the $\eta$s one obtains the following approximation of equation S61

$$p_{obs}(o) = \frac{\int d\eta \, \sum_{t=1}^{T} \delta \left( o - 1 + \frac{1}{T} \sum_{s=1}^{T} \left( \frac{\beta}{\beta + e^{\eta} N_s} \right)^{\beta} \right) \frac{\exp\left( - \frac{-(\eta-\mu)^2}{2\sigma^2} \right)}{\sqrt{2\pi\sigma^2}} \prod_{s=1}^{T} \left( 1 - \left( \frac{\beta}{\beta + e^{\eta} N_s} \right)^{\beta} \right)}{\int d\eta \, \frac{\exp\left( - \frac{-(\eta-\mu)^2}{2\sigma^2} \right)}{\sqrt{2\pi\sigma^2}} \, \prod_{s=1}^{T} \left( 1 - \left( \frac{\beta}{\beta + e^{\eta} N_s} \right)^{\beta} \right)} \tag{S63}$$

Figure S11 shows the empirical occupancy distributions of different biomes, and it compares them with the prediction of equation S61 and with the approximation S63.

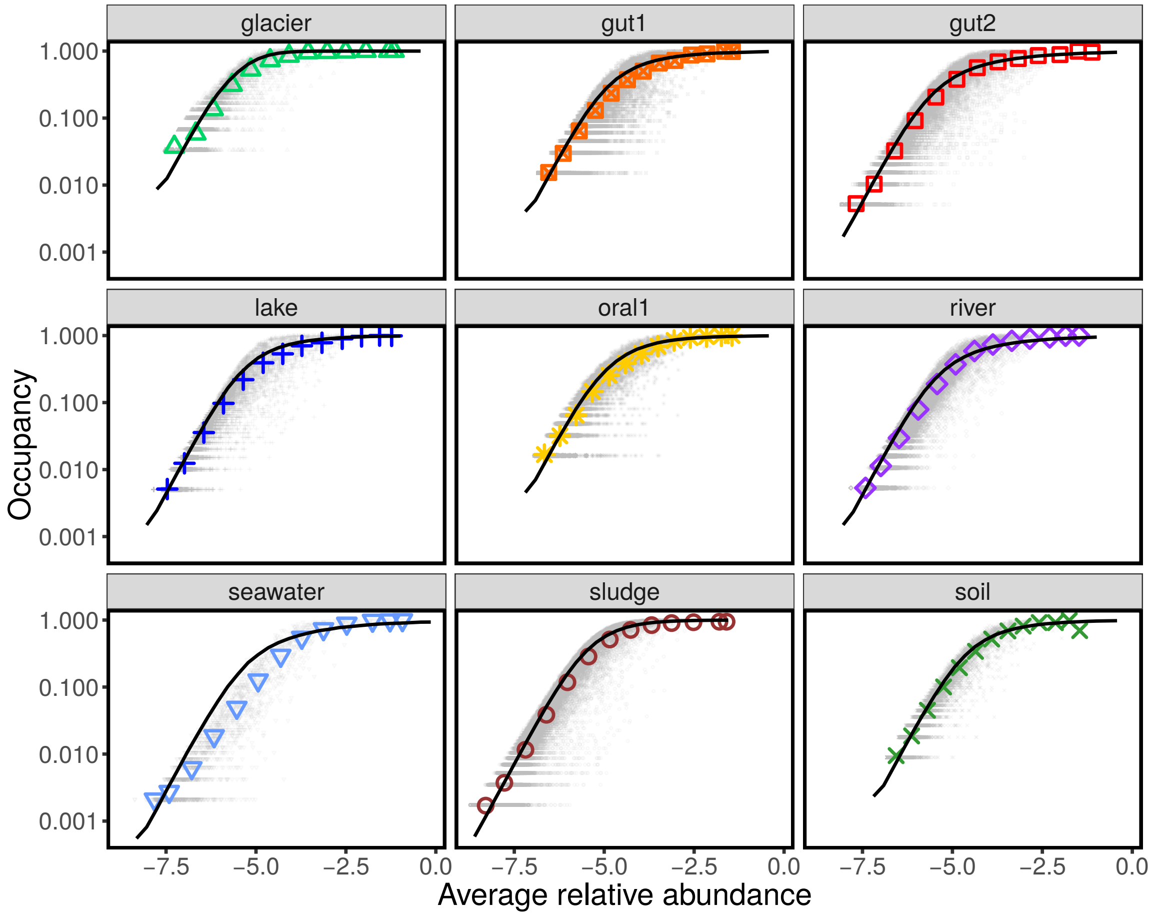


Supplementary Figure S12: **Occupancy-abundance relationship.** The panels report the occupancies $o_i$ vs the log average abundances $\log \bar{x}_i$ of each individual species (gray points). The colored points are averages over species, binned by abundance. The solid black line is the prediction obtained from equation S64.

#### D. Abundance-occupancy relation

Occupancy (the fraction of samples where a species is found) and average abundance are not independent properties. Given an average (relative) abundance $f = \exp(\eta)$, the expected occupancy is

$$\langle o \rangle_\eta = 1 - \frac{1}{T}\sum_{s=1}^{T} P(0|N_s, \eta) = 1 - \frac{1}{T}\sum_{s=1}^{T}\left(\frac{\beta}{\beta + e^\eta N_s}\right)^\beta \,, \tag{S64}$$

where $\eta$ is equal to the logarithm of the average abundance. Figure S21 shows the empirical occupancy-abundance relationship and its comparison with data.

#### E. Species Abundance Distribution

One of the most studied patterns in ecology is the Relative Species Abundance (RSA), which is defined as the number of species with a given abundance. The expected number of species with abundance $n$ in a sample with $N$

total number of reads (i.e., the expected RSA) is given by

$$\langle s_n(N)\rangle = s_{tot}P(n|N) = s_{tot}\int d\eta\ \frac{\Gamma(\beta+n)}{n!\Gamma(\beta)}\left(\frac{e^\eta N}{\beta+e^\eta N}\right)^n\left(\frac{\beta}{\beta+e^\eta N}\right)^\beta \frac{\exp\left(-\frac{-(\eta-\mu)^2}{2\sigma^2}\right)}{\sqrt{2\pi\sigma^2}}\ . \tag{S65}$$

Consistently to equation S57, the number of observed species $\langle s(N)\rangle$ is given by

$$\langle s(N)\rangle = \sum_{n=1}^{\infty}\langle s_n(N)\rangle = s_{tot}\left(1-P(0|N)\right)\ . \tag{S66}$$

In order to compare different samples, it is often more convenient to study the Species Abundance Distribution (SAD), which is defined as the fraction of species with a given abundance. According to our model, the expected SAD is given by

$$\langle \Phi_n(N)\rangle := \frac{\langle s_n(N)\rangle}{\langle s(N)\rangle} = \frac{P(n|N)}{1-P(0,N)} = \frac{\int d\eta\ \frac{\Gamma(\beta+n)}{n!\Gamma(\beta)}\left(\frac{e^\eta N}{\beta+e^\eta N}\right)^n\left(\frac{\beta}{\beta+e^\eta N}\right)^\beta \frac{\exp\left(-\frac{-(\eta-\mu)^2}{2\sigma^2}\right)}{\sqrt{2\pi\sigma^2}}}{1-\int d\eta\ \left(\frac{\beta}{\beta+e^\eta N}\right)^\beta \frac{\exp\left(-\frac{-(\eta-\mu)^2}{2\sigma^2}\right)}{\sqrt{2\pi\sigma^2}}}\ . \tag{S67}$$

The cumulative SAD is defined as

$$\langle \Phi_n^{>}(N)\rangle := \sum_{m=n}^{\infty}\langle \Phi_m(N)\rangle = \frac{\int d\eta\ I_{\frac{e^\eta N}{\beta+e^\eta N}}(n,\beta)\frac{\exp\left(-\frac{-(\eta-\mu)^2}{2\sigma^2}\right)}{\sqrt{2\pi\sigma^2}}}{1-\int d\eta\ \left(\frac{\beta}{\beta+e^\eta N}\right)^\beta \frac{\exp\left(-\frac{-(\eta-\mu)^2}{2\sigma^2}\right)}{\sqrt{2\pi\sigma^2}}}\ , \tag{S68}$$

where $I_p(n,\beta)$ is the regularized incomplete Beta function. Figure S22 compares the empirical cumulative SADs with the prediction of equation S68.

#### S9. MACROECOLOGICAL LAWS IN TEMPORAL DATA

##### A. Law #1: Fluctuations of OTUs abundance across samples are Gamma distributed

Figure S14 shows the distribution of abundance fluctuations across times for the species with high occurrence. This figure parallels Figure S1, which was obtained using the fluctuations across communities instead of across times. Figure S15 shows the estimated moment generating function, obtained in the same way as of Figure S2.

###### *1. Excluding competitive exclusion in time data*

Using the same method explained in section S4 A, we obtained a prediction for the occupancy of each species, based on its average and variance of abundance, which is shown in Figure S16. Also for time data, the presence and absence of species abundances can be predicted from their mean and variance of abundances.

##### B. Law #2: Taylor's law for abundances fluctuations

Figure S17 shows that the quadratic relationship between mean and variance of abundance also hold for time data. These panels parallel Figure S6 obtained across communities.

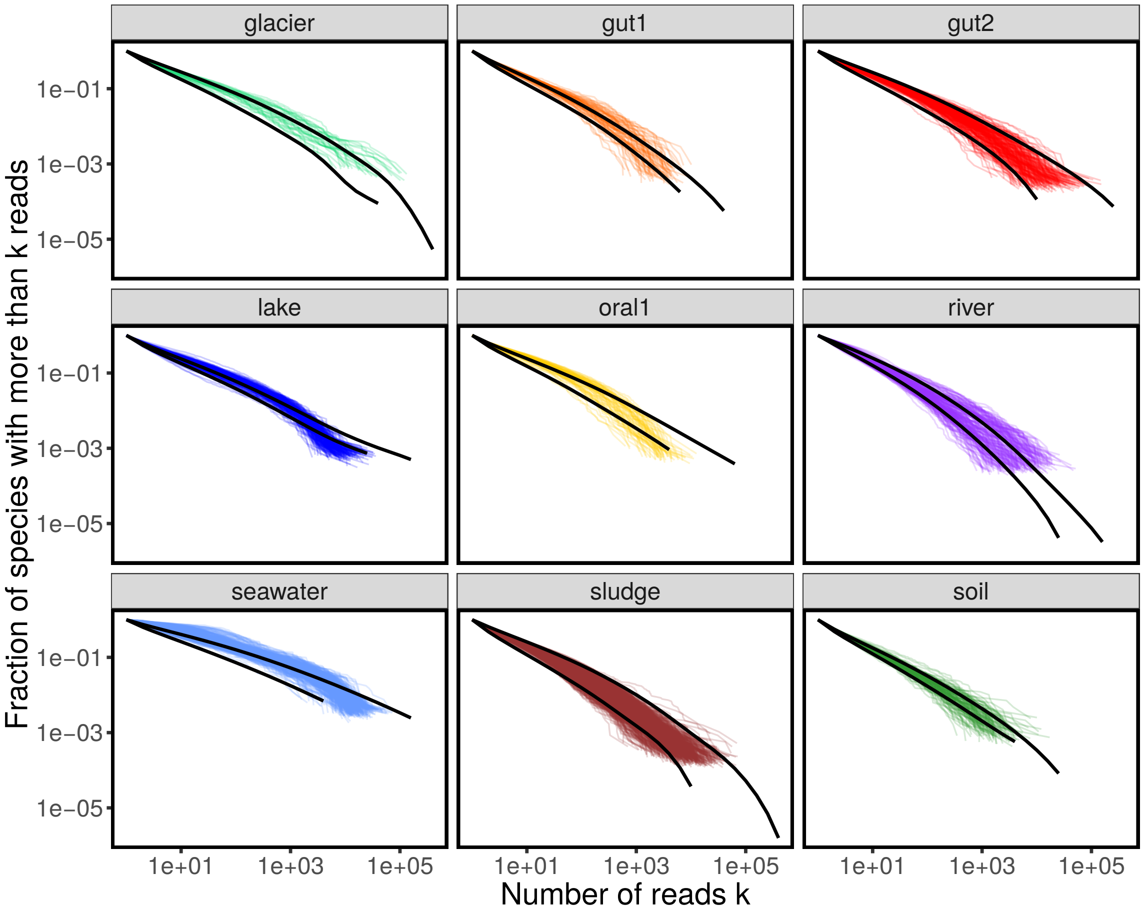


Supplementary Figure S13: **Cumulative Species Abundance Distribution.** The panels report the cumulative Species Abundance Distribution for individual communities (colored lines). The SAD are expected to be influenced by sampling effects, and, in particular, by the total number of reads. The two solid black lines are the expected cumulative SAD for the smallest (on the bottom) and largest (on top) values of total number of reads of each biome.

##### C. Law #3: average abundances are Lognormally distributed

Figure S18 shows that the the average abundances are Lognormally distributed also for time data. These panels parallel Figure S8 obtained across communities.

#### S10. PREDICTION OF MACROECOLOGICAL PATTERNS IN TEMPORAL DATA

As reported in section S8 for cross-sectional data, in this section we show that the three macroecological laws are sufficient to predict other commonly studied macroecological patterns also for temporal data.

Figure S19 shows that equation S57 well predicts the observed number of species as a function of the total number of reads. Figure S20 compares data and the prediction of equation S61 for the occupancy distribution, while Figure S21 shows that equation S64 captures the relationship between average abundance and occupancy. Finally, Figure S22 shows that the prediction for the Species Abundance Distribution obtained in equation S68 in is good agreement with

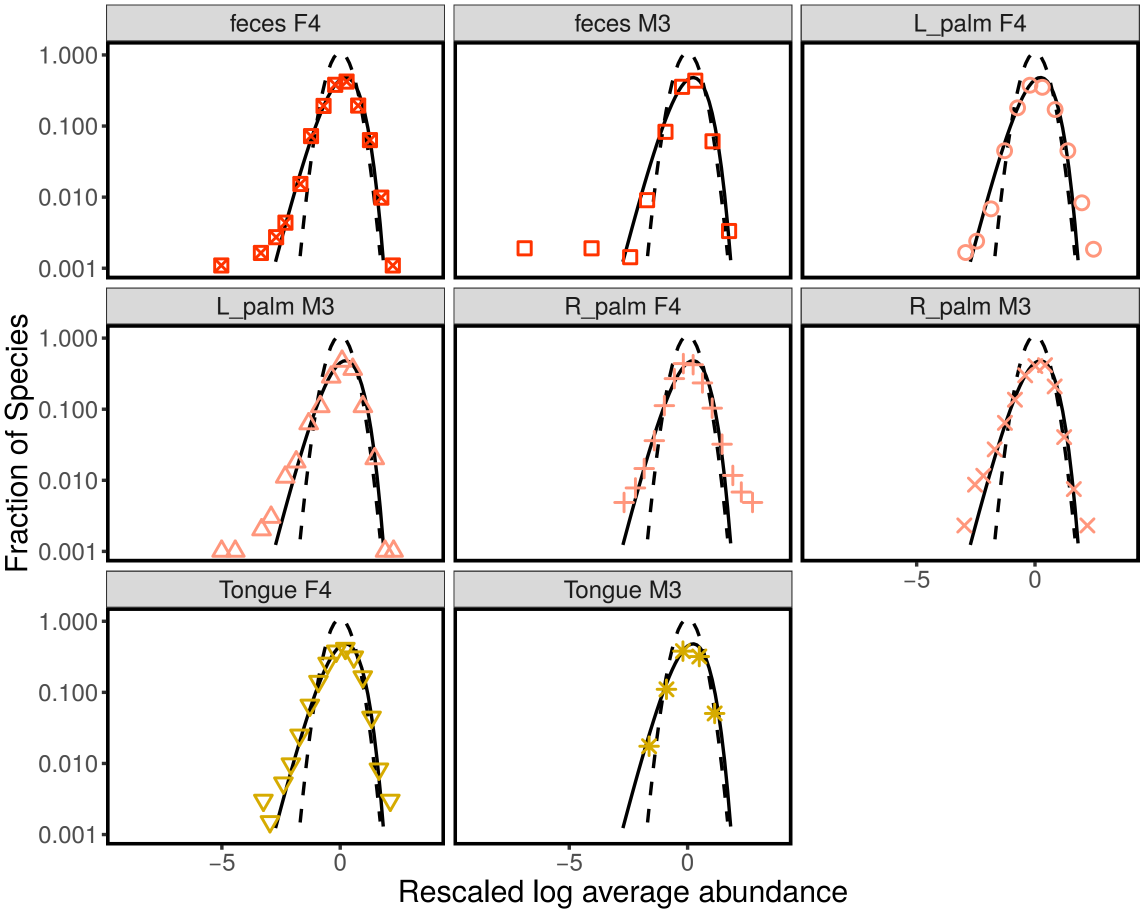


Supplementary Figure S14: **Fluctuations of species abundance over time.** These panels report exactly the same data shown in 4. For each time-series, they were considered only the species present in all the communities. The logarithm of their relative abundances were rescaled (so to have mean zero and unitary variance). The panels report the distribution of these rescaled fluctuations for each biome. Colored points are calculated averaging over both communities and species (same as shown is figure 4). Gray lines are the distribution for individual species over communities. The black continuous line is a Gamma distribution and the black dashed line a Lognormal distribution.

the data.

#### S11. SOURCE OF VARIABILITY IN TEMPORAL DATA

The existence of the same macroecological laws for temporal and cross-sectional data suggests that most of the variability observed across times and across communities has the same origin, that we identified as temporal stochasticity. If the variability was only due to temporal stochasticity, two time snapshot of the same community taken sufficiently apart would be as correlated as the abundance profiles of two different communities. Clearly, we do not claim that this extreme hypothesis is the real scenario. We claim that the departure from this extreme is “small”. The meaning and the precise quantitification of this departure is the focus of this section.

In order to assess this hypothesis, we can compare the correlations of average species abundances within a host and

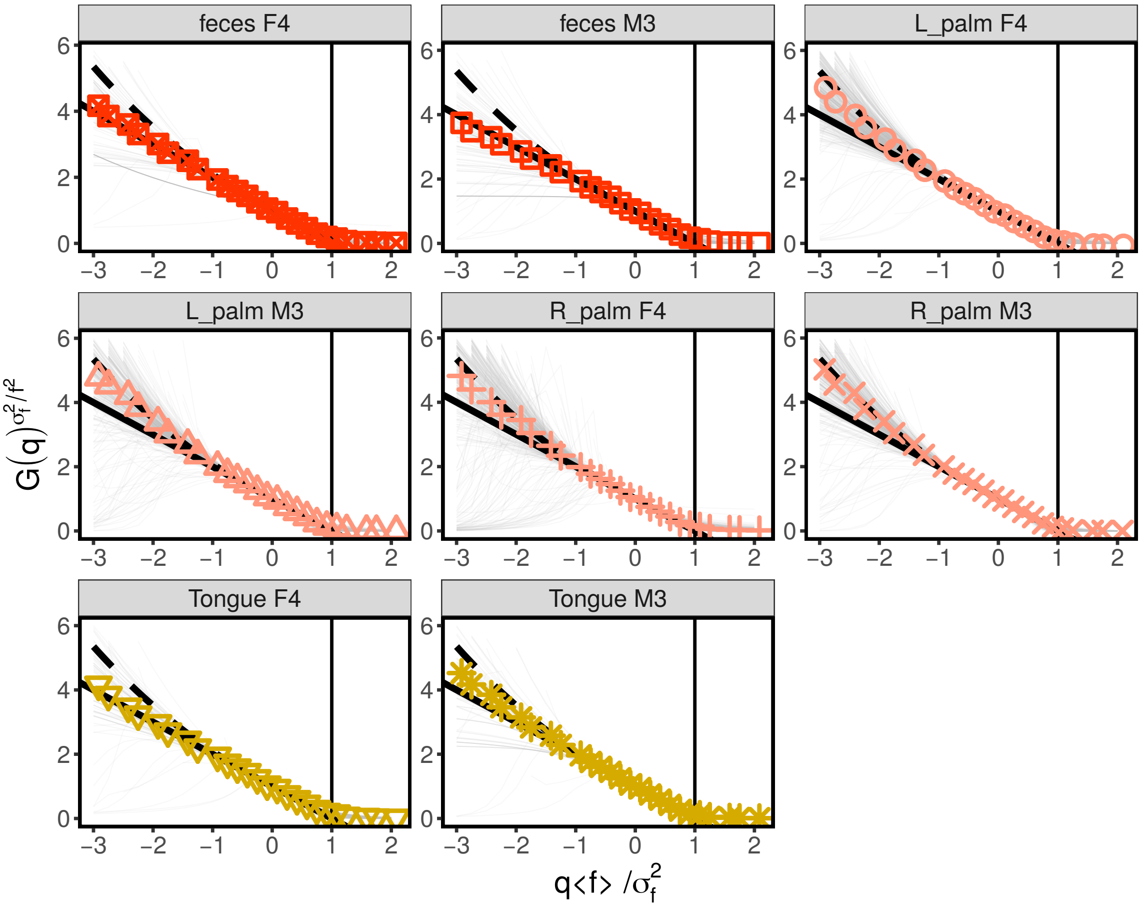


Supplementary Figure S15: **Moment generating function of species fluctuation distribution estimated from time data.** The panels show the moment generating function estimated from the data using equation S32. The gray lines were obtained for individual species (with average average abundance $\bar{x}_i > 5 \cdot 10^{-5}$), while colored points are averages over species. The black solid line is the prediction for the Gamma distribution, while the dashed line is the one for a Lognormal.

between hosts. As a first approximation, the observed correlation between the average abundance in the two samples can be written as

$$c^{ab}_{obs} = c^{ab} \frac{\sigma_x^2}{\sigma_x^2 + \sigma_v^2} \ , \tag{S69}$$

where $c^{ab}$ is the actual correlation, $\sigma_v^2$ is the variance of some noise that act on top of the process and $\sigma_x^2$ is the actual variance of the average abundances. The term $\sigma_v^2$ can be more generally interpreted as any source of variation, indipendent between the two samples, which is affecting the correlation.

In our case we want to estimate how much averages over time differ from average between communities. Let $c_{cross}$ be the correlation of average species abundance across communities and $c_{time}$ the correlation between set of samples obtained at different times. In analogy with S69 we have

$$c_{cross} = c_{time} \frac{\sigma_x^2}{\sigma_x^2 + \sigma_v^2} \ , \tag{S70}$$

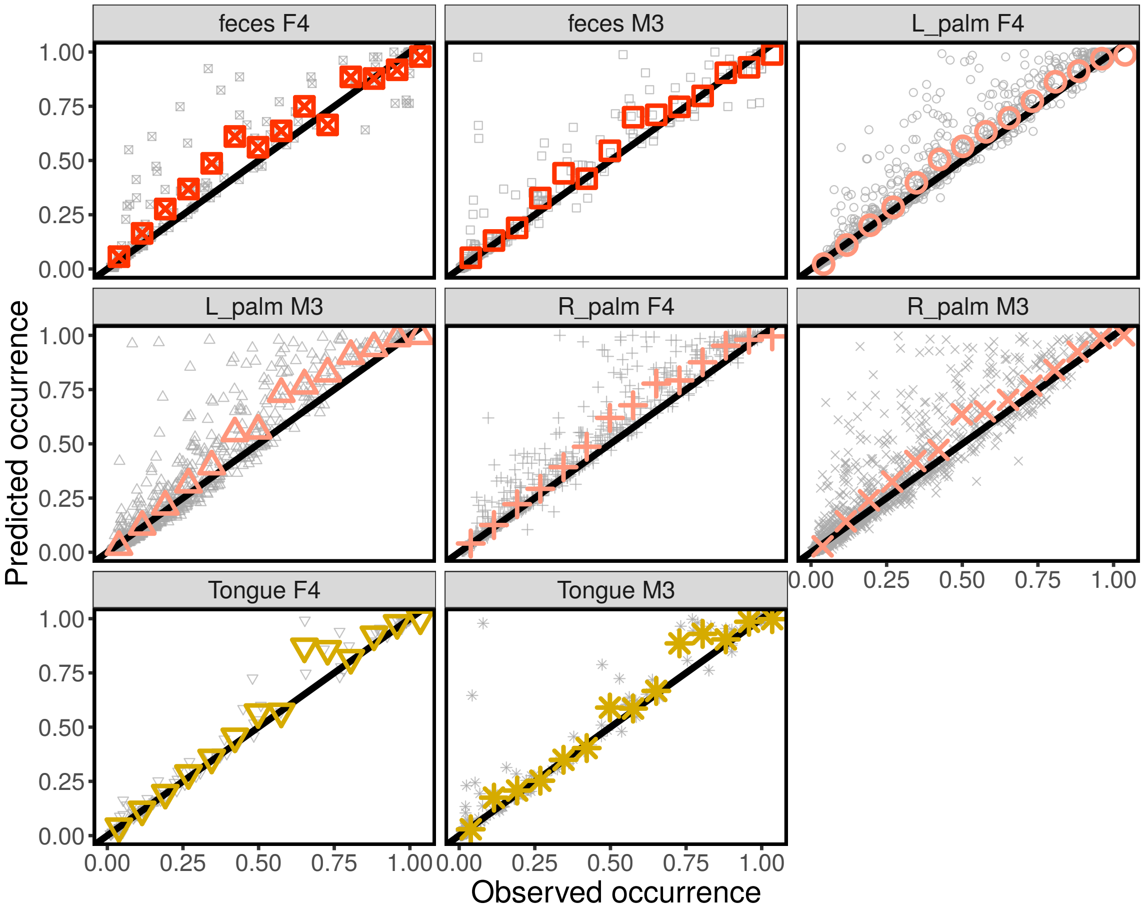


Supplementary Figure S16: **A Gamma AFD correctly predicts species' occupancies for time data.** The occupancy is defined as the fraction of time points where a given species is found to be present. The predicted occupancy was obtained using equation S37, which assumes a Gamma AFD. Using the average and the variance of species' relative abundances, one can in fact estimate the parameters of the AFD and the probability that a species is not found at a given time, given the level of sampling. The black line is the 1 : 1 line, indicating a correct prediction. The gray points are individual species (no filter on average abundance was applied), while the colored points are averages over species.

where, in this context, $\sigma_v/\sigma_x$ is the relative importance of community-specific effects. We obtained therefore that

$$\frac{\sigma_v}{\sigma_x} = \sqrt{\frac{c_{time}}{c_{cross}} - 1}\ , \tag{S71}$$

Table S3 reports the estimate for $\sigma_v/\sigma_x$ using the datasets considered in this work. This relative variation due to community specific effect is extremely small in Oral and Skin communities. The Gut microbiome displays larger variation, but the relative variation due to community specific effect is still fairly small, with a $\frac{\sigma_v}{\sigma_x}$ ratio equal approximately to 0.3.

#### S12. MODELS THAT DO NOT REPRODUCE THE OBSERVED PATTERNS

In this section we compare the the three macroecological laws discussed in section S3, S7, and S5 with the predictions of commonly used theory and models. By design, we use three "laws" that were observed across biomes and therefore

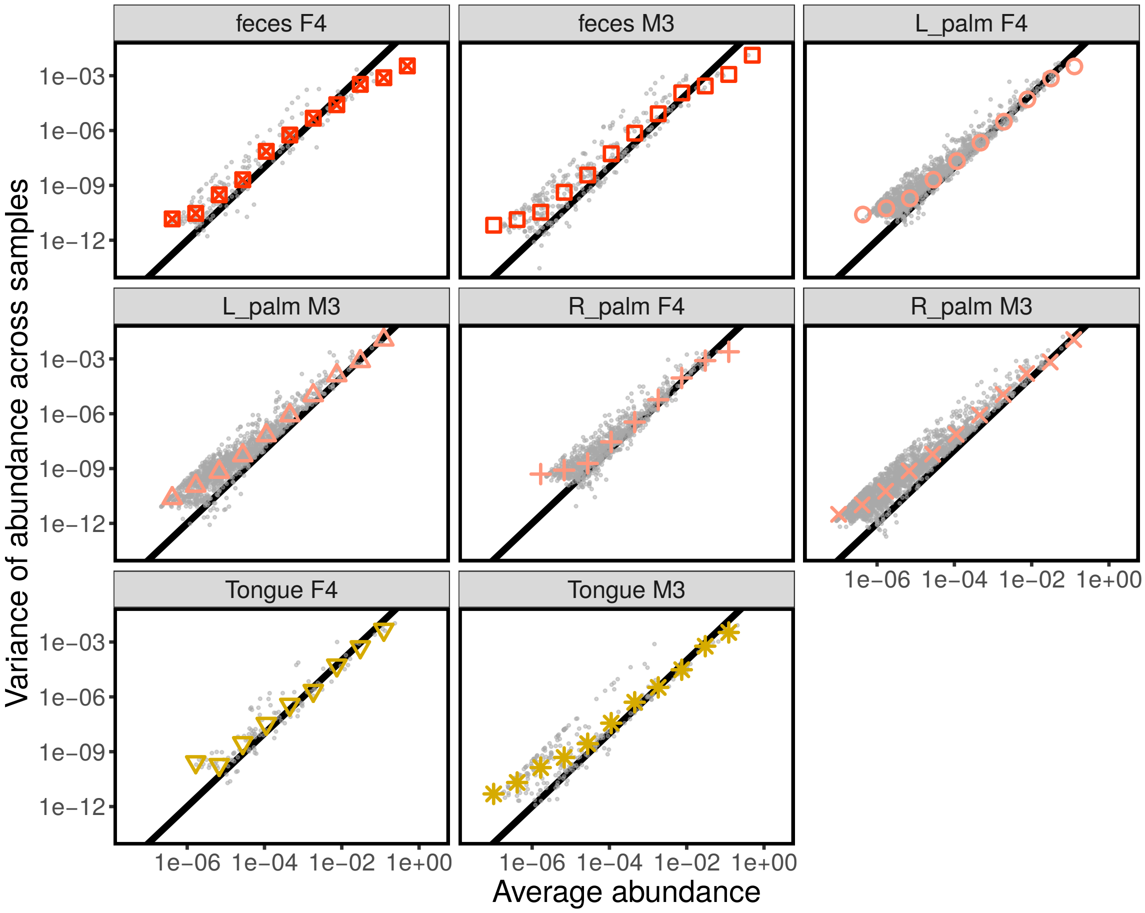


Supplementary Figure S17: **Taylor's law for time data.** The panels show the relationship between mean and variance of abundance. The black line has slope 2, representing the quadratic relationship between variance and mean abundance.

Supplementary Table S3: Estimate of relative cross-community variation.

| Biome ID | $c_{time}$ | $c_{cross}$ | $\sigma_v/\sigma_x$ |
|---|---|---|---|
| Gut | 0.92 | 0.84 | 0.31 |
| Left palm | 0.91 | 0.93 | $\sim 0$ |
| Right Palm | 0.92 | 0.94 | $\sim 0$ |
| Tongue | 0.931 | 0.930 | 0.03 |

they are not specific and cannot reveal too many details about ecological mechanisms that act differently in those biomes. This is also the strength of this test and our observables. For any model or theory aiming at explaining community composition and structure at any level of detail, it is a strong requirement to be able to capture the three minimal laws that we have analyzed in this work. Therefore, while the three laws alone are to general to be useful as positive evidence for any model, they are a strict and general test which can be used to invalidate models and/or theories in their standard formulation.

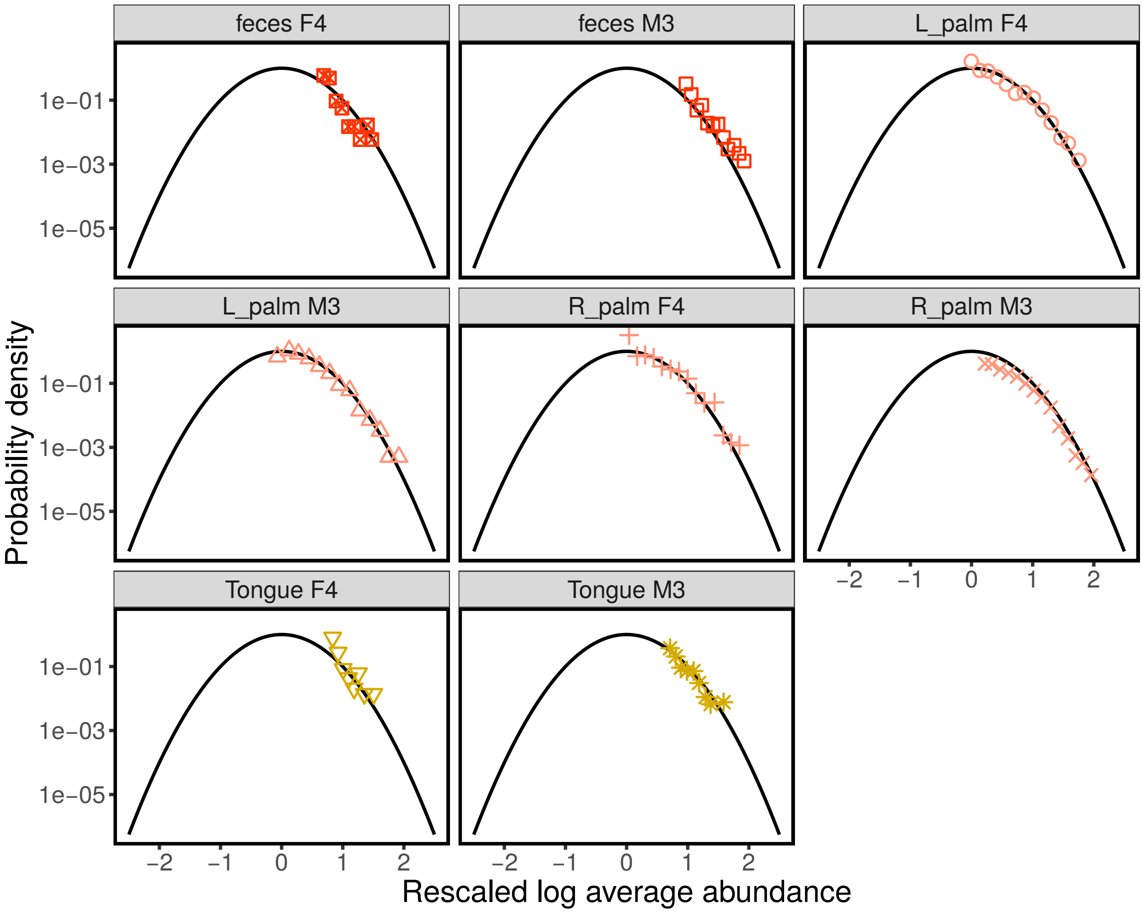


Supplementary Figure S18: **Mean Abundance Distribution (MAD) for time data.** The panels show the distribution of the logarithm of the average abundance $\log \bar{x}_i$. A Lognormal MAD corresponds to normally distributed $\bar{x}_i$. The black line is standardized gaussian distribution (mean zero and variance one). The log average abundances were rescaled as $z_i = (\log \bar{x}_i - \mu)/\sigma$, where $\mu$ and $\sigma$ were obtained for each biome from equation S48. The panels show that the rescaled log average abundances $z_i$ are distributed according to a standard normal, implying that the average abundances $\bar{x}_i$ are lognormally distributed.

##### A. Neutral theory

Neutral theory (NT) [21, 22, 41] has been extremely succesfull in capturing the empirical properties of Relative Species Abundances in many communities [42–44] and it has also proved to be succesfull in capturing dynamical and spatial properties [45]. In the context of microbial communities, NT has been tested in different ways [46]. While the existence of significant correlations (as shown in section S14) contradicts NT, the general patterns of abundance and diversity (and in particular the Relative Species Abundance) do not show a strong disagreement with the prediction of NT. By disentangling the shape of the SAD as the result of the combination of a Gamma AFD and a Lognormal MAD, we are instead able to reject NT on the sole basis of patterns of abundance fluctuations. In fact, NT would predict a peaked MAD (Gaussian for a finite number of samples), while we observe a Lognormal distribution in the data. Moreover the average abundance is a conserved property, characteristic of some broad environmental conditions (see section S6). This result strongly suggests not only that NT is strictly rejected from the data, but also, more

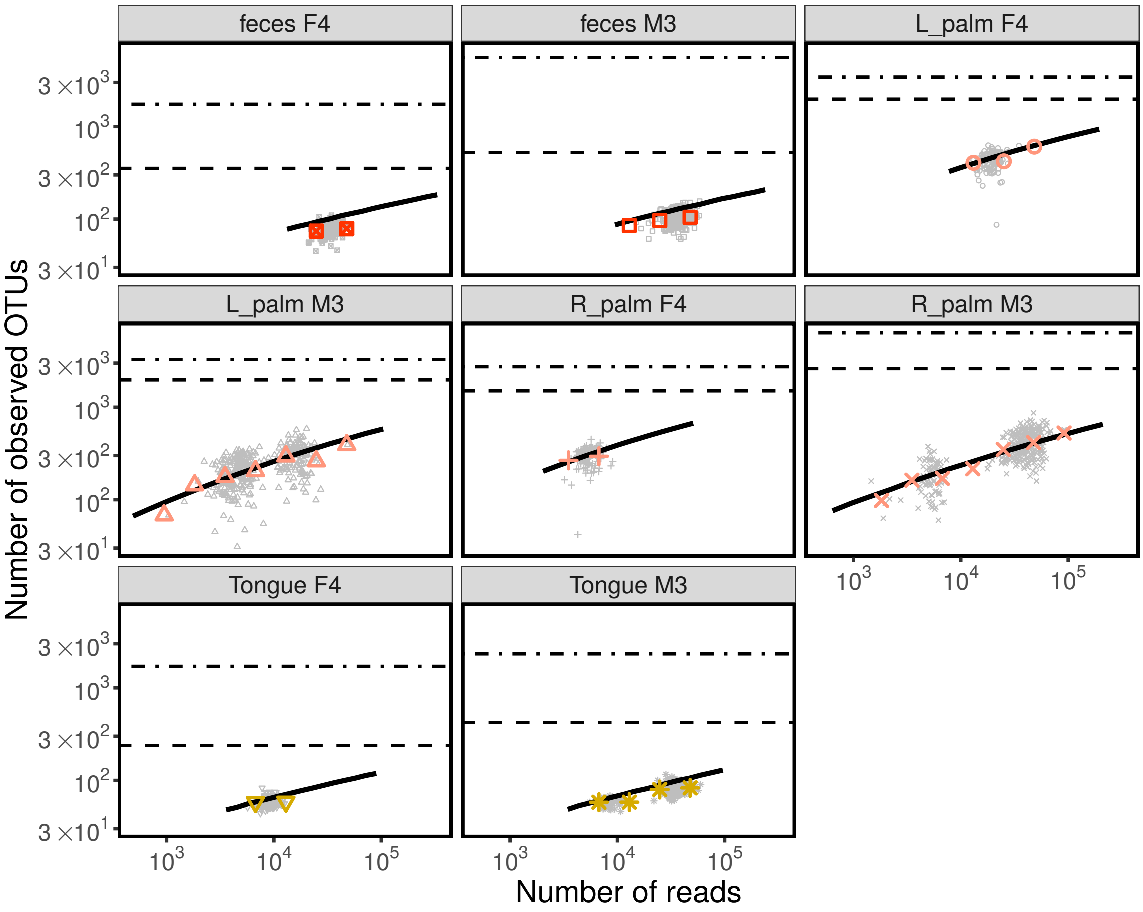


Supplementary Figure S19: **Number of observed species vs. total number of reads for time data.** The number of observed species at a given time, depends on the total number of sequences sampled $N_s$. The more sequences are sampled, the more likely it is to find new species. The gray points report the number of species observed in each community, while the colored symbols are averaged over times in a range of $N_s$. The black dashed line represent $s_{obs}$, the total number of observed species across all the times in that biome. The black dot-dashed line reports the value of the inferred value of $s_{tot}$, which includes also the species that have not been observed in that biome. The solid black line is the prediction, reported in equation S57, obtained by combining the three macroecological laws. Equation S57 correctly predicts the typical values of observed number of species, as well as the quantitative relationship between species and number of sequences.

importantly, that it is not the right starting point to formulate more realistic models on species presence and abundance variation.

##### B. Deterministic models with alternative stable states as source of variation

Lotka-Volterra [47, 48] or consumer-resource models [49] often leads to alternative stable states driven by competitive exclusion: only a subset of the species can coexist in the same community. Which subset of species is realized in a community depends on the initial condition and on its basin of attraction. Within a biome (whose definition is unclear, a priori) we showed that the presence/absence of species is a consequence of sampling errors. This also applies in a single host when observed over time. While abundance of species fluctuates, the average value around which it

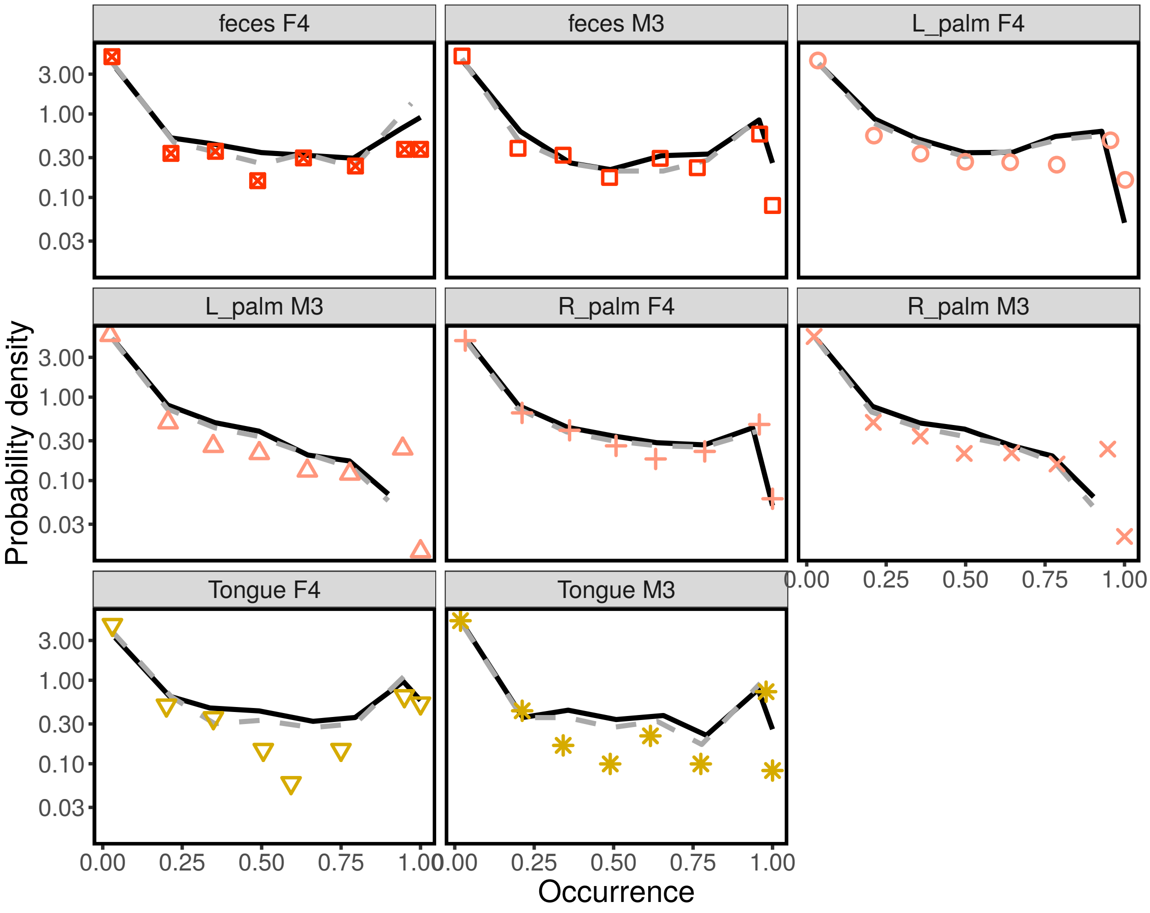


Supplementary Figure S20: **Distribution of occupancy of species across times.** The colored symbols report the distribution of species' occupancies (the fraction of times when a species has been observed) in each biome. The solid black line is the prediction, reported equation S61, obtained by combining the three macroecological laws. The gray dashed line is the approximation to the prediction reported in equation S63.

fluctuates is conserved over time and across communities. This observations suggest that, as a first approximation, it exists in fact only one basin of attraction (or some globally stable equilibrium) around which species fluctuates. The sentence "as a first approximation" should be interpreted as to characterize the main contributors of fluctuations from community to community, from sample to sample, which are very likely not to be due to alternative stable states.

#### S13. STOCHASTIC LOGISTIC EQUATION

We assume that the dynamics of population abundance $x_i$ is defined by

$$\frac{dx_i(t)}{dt} = \frac{\dot{x}_i(t)}{\tau_i}\left(1 - \frac{x_i(t)}{K_i}\right) + \sqrt{\frac{\sigma_i}{\tau_i}}x_i(t)\xi_i(t) \ , \tag{S72}$$

where $\xi_i(t)$ is a Gaussian delta correlated white noise ($\langle \xi_i(t) \rangle = 0$ and $\langle \xi_i(t)\xi_j(t') \rangle = \delta_{ij}\delta(t - t')$).

The term proportional to $\xi_i(t)$ represents environmental fluctuations, which translate into fluctuations of the growth rate and are therefore proportional to $x_i$. The only essential assumption here is that we assume these fluctuations to

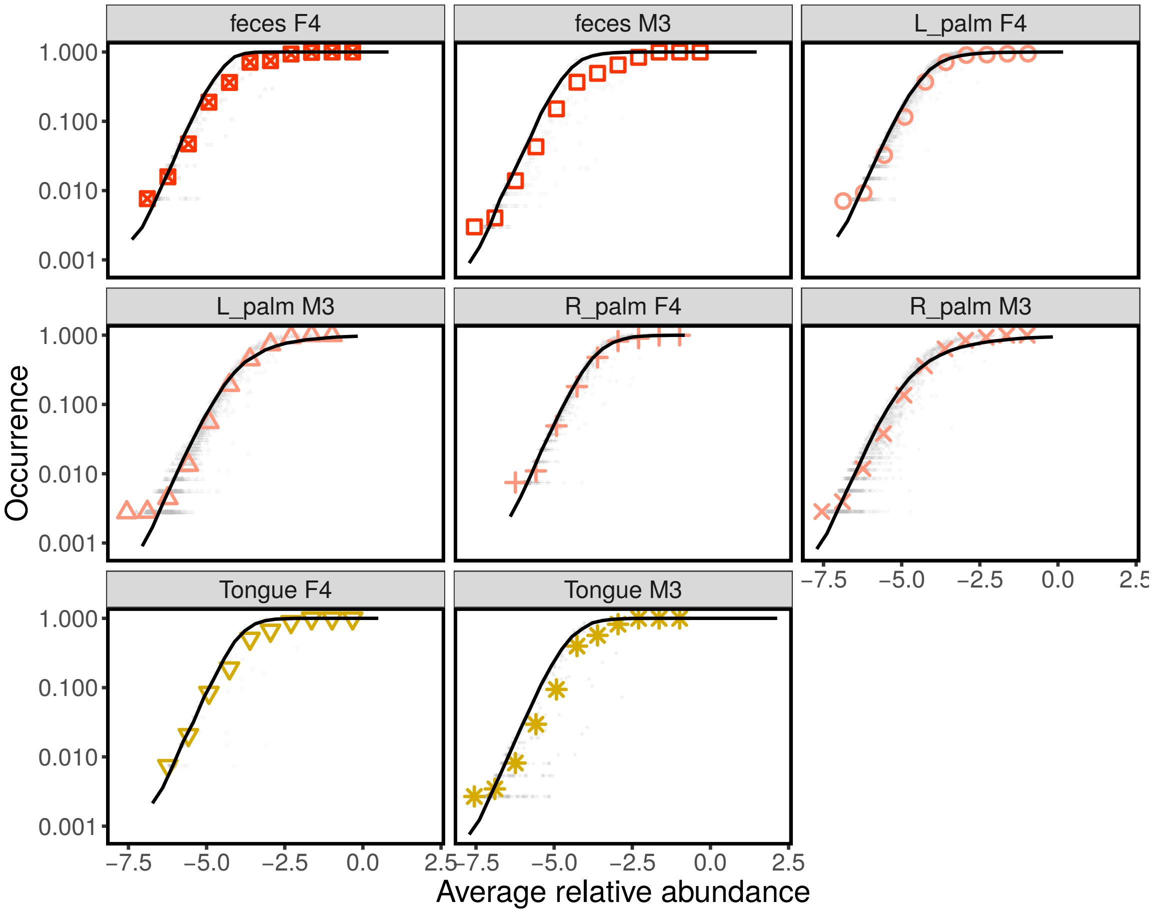


Supplementary Figure S21: **Occupancy-abundance relationship for time data.** The panels report the occupancies $o_i$ vs the log average abundances $\log \bar{x}_i$ of each individual species (gray points). The colored points are averages over species, binned by abundance. The solid black line is the prediction obtained from equation S64.

occurr at a typical timescale that is much shorter than the population dynamics one and to have a finite variance. From this assumption it follows that $\xi_i(t)$ can be well approximated as a Gaussian delta correlated white noise. The other term is a logistic growth term, with carrying capacity $K_i$. The parameter $\tau_i$ is the timescale of population dynamics and $\sigma_i$ is measures the coefficient of variation of growth rate fluctuations.

Interpreted with Itô perscription [50], equation S72 corresponds to the following Fokker-Planck equation which describes the dynamics of the probability $P_i(x,t)$ of finding species $i$ with abundance $x$ at time $t$

$$\tau_i \frac{\partial P_i(x,t)}{\partial t} = -\frac{\partial}{\partial x}\left( x\left(1 - \frac{x}{K_i}\right) P_i(x,t)\right) + \frac{\sigma_i}{2}\frac{\partial^2}{\partial x^2}\left(x^2 P_i(x,t)\right) \ . \tag{S73}$$

The stationary distribution $P_i^*(x) = \lim_{t\to\infty} P(x,t)$ can be found by setting the left hand side of equation S73 equal to zero. Imposing detailed balance, we obtain

$$x\left(1 - \frac{x}{K_i}\right) P_i^*(x) = \frac{\sigma_i}{2}\frac{\partial}{\partial x}\left(x^2 P_i^*(x)\right) \ . \tag{S74}$$

By introducing $x^2 P_i^*(x) = Q_i(x)$ we obtain

$$\left(\frac{1}{x} - \frac{1}{K_i}\right) Q_i(x) = \frac{\sigma_i}{2} Q_i'(x) \ , \tag{S75}$$

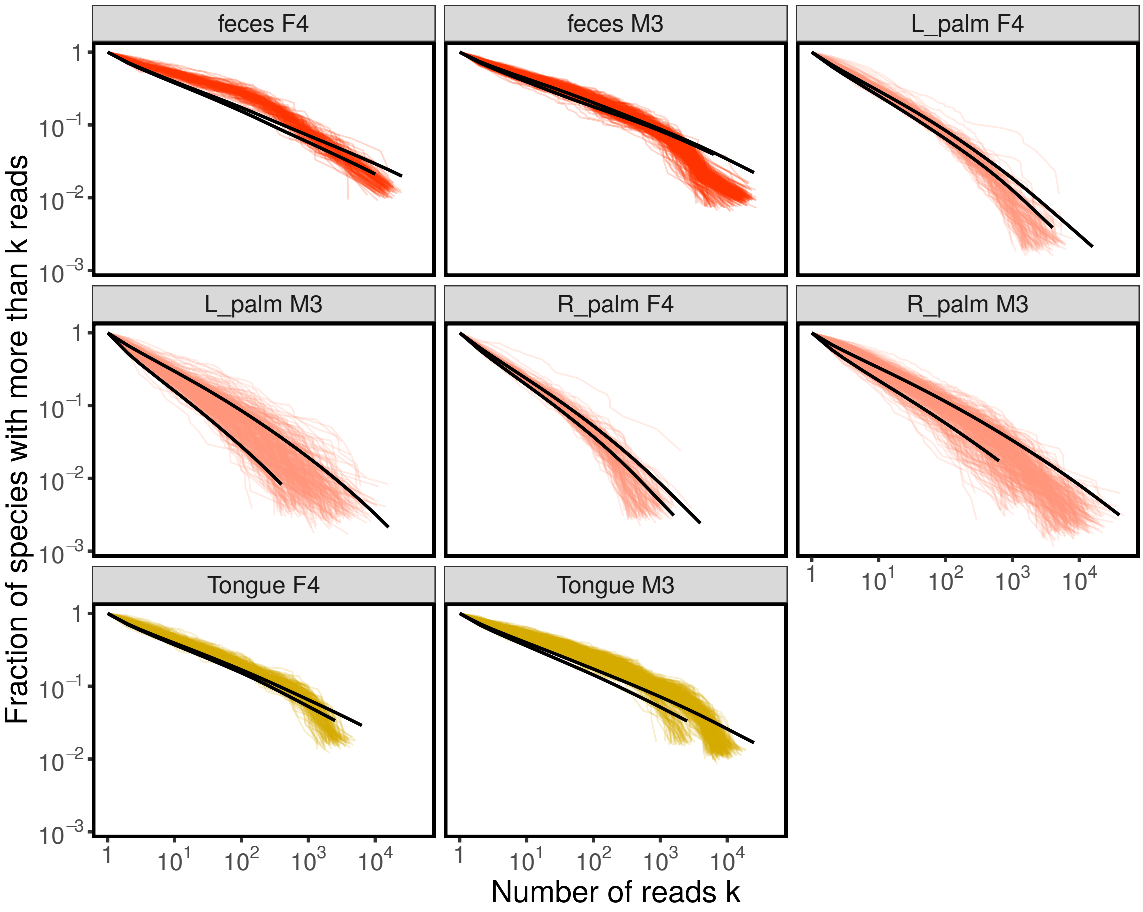


Supplementary Figure S22: **Cumulative Species Abundance Distribution for time data.** The panels report the cumulative Species Abundance Distribution for individual times (colored lines). The SAD are expected to be influenced by sampling effects, and, in particular, by the total number of reads. The two solid black lines are the expected cumulative SAD for the smallest (on the bottom) and largest (on top) values of total number of reads of each biome, obtained from equation S68.

which is solved by

$$Q_i(x) = c \exp\left(-\frac{2}{K_i \sigma_i} x\right) x^{2\sigma^{-1}} \,, \tag{S76}$$

where $c$ is an arbitrary constant. By using $P_i^*(x) = Q_i(x) x^{-2}$ and fixing $c$ by imposing $\int dx P_i^*(x)$, we obtain

$$P_i^*(x) = \frac{1}{\Gamma(2\sigma_i^{-1} - 1)} \left(\frac{2}{K_i \sigma_i}\right)^{2\sigma_i^{-1} - 1} \exp\left(-\frac{2}{K_i \sigma_i} x\right) x^{2\sigma_i^{-1} - 2} \,, \tag{S77}$$

which is a Gamma distribution with mean

$$\langle x_i \rangle = K_i (1 - \frac{\sigma}{2}) \,, \tag{S78}$$

and coefficient of variation

$$\frac{\langle x_i^2 \rangle - \langle x_i \rangle^2}{\langle x_i \rangle^2} = \frac{\sigma}{2 - \sigma} \,. \tag{S79}$$

It is important to observe that the mean abundance is positive, and the variance is finite only if $\sigma < 2$. If the environmental fluctuations are too strong the population is, in fact, driven to exctintion. This effect is driven by the multiplicative nature of the noise in equation S73. If we introduce the variable $q_i = \log x_i$ in equation S73, we obtain (under Itô perscritpion)

$$\frac{dq_i(t)}{dt} = \frac{1}{\tau_i}\left(1 - \frac{\sigma}{2} - \frac{e^{q_i}}{K_i}\right) + \sqrt{\frac{\sigma_i}{\tau_i}}\xi_i(t) \ . \tag{S80}$$

By taking the average on both sides we obtain

$$\frac{d\langle q_i(t)\rangle}{dt} = \frac{1}{\tau_i}\left(1 - \frac{\sigma}{2} - \frac{\langle e^{q_i}\rangle}{K_i}\right) \ . \tag{S81}$$

Since the term $\langle e^{q_i}\rangle$ is always positive, it is evident that $\frac{d\langle q_i(t)\rangle}{dt}$ reaches a stationary value only if $1 - \sigma/2 > 0$, i.e. if $\sigma < 2$. If $\sigma > 2$ the population abundance decreases indefinitely.

#### S14. CORRELATIONS OF SPECIES ABUNDANCE FLUCTUATIONS

In the previous sections we showed that the three macroecological laws correctly capture many statistical properties of the empirical data. In this section we show that they do not capture all the statistical properties, and, in particular, they fail in describing the correlations between species abundance fluctuations.

In section S8, in order to reproduce the macroecological patterns starting from the three macroecological laws, we assumed that the abundance fluctuations were independent across. More generally we can write

$$P_{ij}(n_i, n_j|N) = \int dx\ dy\ \frac{(xN)^{n_i}}{n_i!}e^{-xN}\ \frac{(yN)^{n_i}}{n_i!}\frac{(yN)^{n_i}}{n_i!}e^{-yN}\ \rho_{ij}(x,y) \ , \tag{S82}$$

where $P_{ij}(n_i, n_j|N)$ is the probability of observing $n_i$ reads of species $i$ and $n_j$ reads of species $j$ in a sample with $N$ total number of reads, and $\rho_{ij}(x,y)$ is the joint probability distribution of the (relative) abundances. So far we have assumed $\rho_{ij}(x,y) = \rho_i(x)\rho_j(y)$ which implies in turn $P_{ij}(n_i, n_j|N) = P_i(n_i, N)P_j(n_j|N)$. Using the same concepts introduced in section S2, it is easy to show that

$$\int dx\ dy\ xy\ \rho_{ij}(x,y) = \langle x_i x_j\rangle \approx \frac{1}{T}\sum_{s=1}^{T}\frac{n_i^s}{N_s}\frac{n_j^s}{N_s} \ , \tag{S83}$$

for a large number of samples $T$. By considering also mean and variance of the two marginal distributions $\rho_i(x)$ and $\rho_j(y)$ we can easily estimate the Pearson correlation coefficient $r_{ij}$. In the limit of large numer of samples, if the abundances were independent, the Pearson correlation coefficient would tend to zero. Since we are working with a finite number of samples, the fluctuations in the estimated correlation coefficient cannot be neglected. Instead of studying all the correlations independently, we considered the distribution of coefficients $r_{ij}$, formally defined as

$$q(r) := \frac{2}{s(s-1)}\sum_{i>j}\delta(r - r_{ij}) \ , \tag{S84}$$

where $s$ is the number of species considered.

Figure S23 compares the empirical distribution of Pearson correlations $q(r)$, with the one obtained by imposing independence between species and using the three macroecological laws. The first important observation is that the two distibution differs: there are in fact significal correlation which cannot be neglected. The second important observation is that the empirical $q(r)$ is centered about zero: with most of the species pairs have low / non-significal correlation. This second observation agrees qualitatively with the null expectation, but is far from trivial. In fact, it

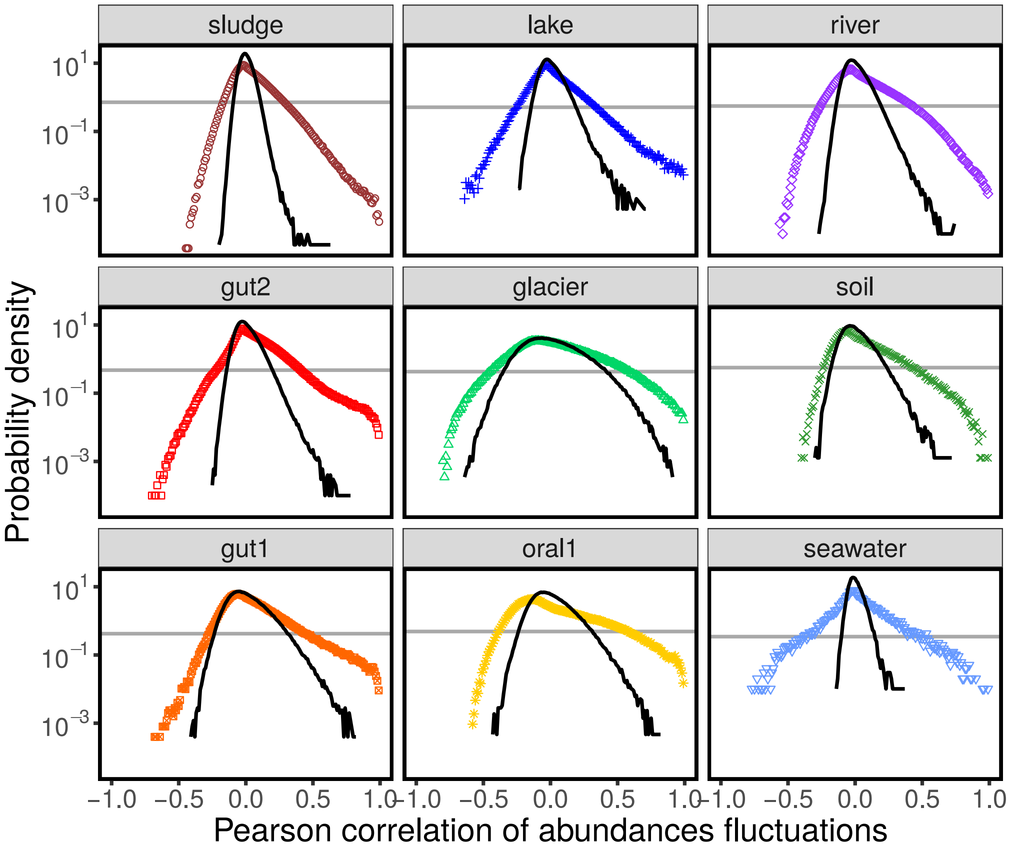


Supplementary Figure S23: **Distribution of Pearson correlation coefficients of abundances fluctuations.** For each biome and each pair of species (present in ar least 50% of the samples), we computed the Pearson correlation of their abundances fluctuations. Each panel shows the distribution of these correlation coefficients (over all the species pairs) for each biome. The colored points are data, while the black line is the null expectation obtained using the first macroecological law, where we empirically fixed the free parameters (empirical mean and variance of each species). They gray horizontal line represent the 95% threshold. These figures convey two important messages: 1) there exist significant correlation which are not captured by the the three macroecological laws alone; 2) correlations are present, but they are weak/sparse: most of pairs of species do not have large correlations.

implies that correlations are weak (correlation coefficient are small) and sparse (pairs of species with large correlations are rare).

These two important results are retrospectively important to interpret the main results of the paper. It is possible to make predictions about the macroecological patterns by assuming only the three macroecological laws and independence between species because correlations are weak, and therefore they do not affect much the macroecological patters (which are averages over species or samples). In turn, the results of the papers are essential to detect correlation, providing an empirically-validated null model.

[10] Brown, J. H. *Macroecology* (University of Chicago Press, 1995).

[21] Hubbell, S. P. *The Unified Neutral Theory of Biodiversity and Biogeography* (Princeton University Press, 2001).

[27] Roy, F., Biroli, G., Bunin, G. & Cammarota, C. Numerical implementation of dynamical mean field theory for disordered systems: application to the Lotka-Volterra model of ecosystems (2019). 1901.10036.

[28] Mitchell, A. L. *et al.* EBI Metagenomics in 2017: enriching the analysis of microbial communities, from sequence reads to assemblies. *Nucleic Acids Research* **46**, D726–D735 (2018).

[29] Gloor, G. B., Macklaim, J. M., Pawlowsky-Glahn, V. & Egozcue, J. J. Microbiome Datasets Are Compositional: And This Is Not Optional. *Frontiers in Microbiology* **8**, 2224 (2017).

[30] Koonin, E. V. *The logic of chance : the nature and origin of biological evolution* (Pearson Education, 2012).

[50] Note that the corresponding Stranovich equation has the same form up to a redefinition of parameters.
